# PrrA modulates *Mycobacterium tuberculosis* response to multiple environmental cues and is critically regulated by serine/threonine protein kinases

**DOI:** 10.1101/2022.03.20.485039

**Authors:** David Giacalone, Rochelle E. Yap, Alwyn M. V. Ecker, Shumin Tan

## Abstract

The ability of *Mycobacterium tuberculosis* (Mtb) to adapt to its surrounding environment is critical for the bacterium to successfully colonize its host. Transcriptional changes are a vital mechanism by which Mtb responds to key environmental signals experienced, such as pH, chloride (Cl^-^), nitric oxide (NO), and hypoxia. However, much remains unknown regarding how Mtb coordinates its response to the disparate signals seen during infection. Utilizing a transcription factor (TF) overexpression plasmid library in combination with a pH/Cl^-^-responsive luciferase reporter, we identified the essential TF, PrrA, part of the PrrAB two-component system, as a TF involved in modulation of Mtb response to pH and Cl^-^. Further studies revealed that PrrA also affected Mtb response to NO and hypoxia, with *prrA* overexpression dampening induction of NO and hypoxia-responsive genes. PrrA is phosphorylated not just by its cognate sensor histidine kinase PrrB, but also by serine/threonine protein kinases (STPKs) at a second distinct site. Strikingly, a STPK phosphoablative PrrA variant was significantly dampened in its response to NO versus wild type Mtb, disrupted in its ability to adaptively enter a non-replicative state upon extended NO exposure, and attenuated for *in vivo* colonization. Together, our results reveal PrrA as an important regulator of Mtb response to multiple environmental signals, and uncover a critical role of STPK regulation of PrrA in its function.

## INTRODUCTION

The remarkable ability of *Mycobacterium tuberculosis* (Mtb) to survive within a host is predicated on its ability to sense and respond to changing environmental signals encountered during infection. These include cues such as pH, chloride (Cl^-^), nitric oxide (NO), and hypoxia[1–4], and we have previously shown that Cl^-^ concentration ([Cl^-^]) increases within the phagosomal compartment during phagosome maturation, simultaneous to acidification, demonstrating the linked nature of these cues [3]. Strikingly, the transcriptional response of Mtb to acidic pH and high [Cl^-^] is synergistic [3]. Similarly, Mtb response to NO and hypoxia is intimately linked, with extensive overlap in the bacterial transcriptional response to these two signals, and the DosRS(T) two-component system (TCS) regulating a core subset of genes in response to either signal [2, 4–7]. These environmental signals change in predictable manners within the host, reflecting both location and the status of the immune response [1, 3, 8–10]. They further affect Mtb growth status, with Mtb growth slowed at acidic pH [11, 12], and the bacteria entering an adaptive state of growth arrest upon extended exposure to NO or hypoxia [2, 13, 14]. The feature of slowed growth at acidic pH is unique to bacteria within the Mtb complex, as mycobacteria not within the Mtb complex do not reduce their growth rate under acidic pH conditions [12, 15]. This intriguingly suggests genetic “hard-wiring” of regulation of Mtb growth in response to environmental cues, and indeed Mtb growth at acidic pH can be manipulated both genetically and via changing of the provided carbon source [16, 17]. Critically, a failure to respond to environmental cues such as pH and NO results in attenuation of the ability of Mtb to colonize its host [18–22]. While studies of Mtb response to environmental cues have largely focused on one signal at a time, Mtb is exposed to multiple, disparate signals simultaneously *in vivo*, and coordination of an adaptive response is essential for the bacterium’s successful colonization of its host.

Transcriptional regulation is a key aspect of Mtb adaptation to the host and response to changes in its local microenvironment, and the Mtb genome encodes 214 annotated or known transcription factors (TFs) [23, 24]. While the role of most of these TFs remain unknown, a few crucial TCSs that regulate Mtb environmental response have been well-characterized and clearly shown to be critical for successful host colonization [19, 22, 25–27]. A key example is the PhoPR TCS that is required for Mtb response to pH and also plays a role in regulation of Mtb response to Cl^-^ [1, 3, 11]. Of note, although a Δ*phoPR* mutant fails to respond to acidic pH, it is still capable of synergistically responding to acidic pH and high [Cl^-^] [3]. Similarly, while the DosRS(T) TCS is known to be vital in regulating Mtb response to NO and hypoxia [2, 4, 5], the DosR regulon consists of a subset of the >200 genes differentially expressed upon Mtb exposure to NO or hypoxia [2, 4]. Together, these data highlight the likely existence of other TFs that play critical roles in regulating Mtb response to these key environmental cues.

An additional layer of Mtb environmental response regulation distinct from transcriptional control is via the action of serine/threonine protein kinases (STPKs) [28, 29]. While STPKs are far less abundant in bacteria as compared to TCSs [30], they are prominent in Mtb, with an almost equivalent number of STPKs (11) present as TCSs (12) [29]. Although hundreds of proteins are predicted to be phosphorylated by STPKs in Mtb, the role of STPKs in Mtb environmental sensing/response remains poorly understood. Nonetheless, more recent genome-wide studies have predicted that many TCSs (9/12) are phosphorylated by STPKs [28, 31], raising the intriguing possibility of an important role for STPK-TCS interplay in Mtb environmental cue response. In particular, phosphorylation of the TCS response regulator DosR by the STPK PknH has been shown biochemically, with a Δ*pknH* Mtb mutant exhibiting impaired NO-mediated induction of DosR regulon genes [32]. Additionally, another study showed that binding to target DNA by the essential response regulator MtrA was decreased in a STPK-phosphomimetic MtrA variant versus wild type MtrA [33]. Importantly, STPK phosphorylation represents a rapid means for altering the activity of a TCS response regulator, separate from slower transcription changes, a possible distinct advantage in the context of quickly changing environmental signals that may be encountered by Mtb during infection.

In this study, we exploit a pH/Cl^-^-responsive luminescent reporter Mtb strain in combination with a tetracycline-inducible TF overexpression plasmid library in a phenotypic screen, identifying PrrA, an essential TF that is part of the PrrAB TCS, as a global modulator of Mtb response to not just pH and Cl^-^, but also NO and hypoxia. Utilizing an inducible overexpression system, we show that *prrA* levels are critical for proper regulation of Mtb response to these environmental cues. Strikingly, we discover that STPK phosphorylation of PrrA plays a vital role in regulating PrrA function, with a STPK-phosphoablative PrrA Mtb variant significantly disrupted in environmental cue response, ability to enter an adaptive state of growth arrest upon extended exposure to NO, and host colonization.

## RESULTS

### Exploitation of a Cl*^-^*/pH responsive reporter strain in combination with a transcription factor (TF) overexpression library uncovers a role for PrrA in Mtb response to pH and Cl*^-^*

To identify TFs that modulate the response of Mtb to pH and Cl^-^, we first constructed a luciferase version of our previously established Cl^-^/pH-responsive *rv2390c’*::GFP reporter [3] by fusing the promoter region of *rv2390c* to a gene encoding firefly luciferase, and integrating this luminescent reporter into the chromosome of Mtb (*rv2390c’*::luciferase). As expected, robust upregulation in light output was observed in acidic pH (pH 5.7) or high Cl^-^ concentration ([Cl^-^], 250 mM NaCl) conditions, with a synergistic response in the dual condition of pH 5.7 + 250 mM NaCl, consistent with previous observations using the GFP reporter (Fig. 1A) [3]. An arrayed library of inducible TF overexpression strains carrying the *rv2390c’*::luciferase reporter was then generated by individually transforming each of 207 tetracycline-inducible TF overexpression plasmids [23, 24] into the *rv2390c’*::luciferase reporter Mtb strain. The library was screened in (i) 7H9, pH 7.0 (control), (ii) 7H9, pH 5.7, (iii) 7H9, pH 7.0, 250 mM NaCl, or (iv) 7H9, pH 5.7, 250 mM NaCl conditions, with utilization of an arrayed format to eliminate bottleneck effects and enable immediate identification of any hits of interest from the screens. Identification of *phoP* and *sigE* as hits where overexpression increased *rv2390c’*::luciferase reporter signal compared to the empty vector control in the high [Cl^-^] (250 mM NaCl) screen condition (8.5 and 25-fold respectively versus an average of 4.3-fold induction in the empty vector controls, with fold induction calculated as relative light units (RLU)/OD_600_ in the test versus control condition to account for any differences in bacterial growth) served as validation that the screen functioned as anticipated, as a Δ*phoP* mutant strain has previously been shown to dampen Mtb response to Cl^-^, while SigE has been shown to interact with PhoP in the regulation of Mtb response to acidic pH (Fig. 1B, Table S1) [3, 34]. Both of these genes were also observed as hits that increased *rv2390c’*::luciferase reporter signal in the acidic pH (pH 5.7) and acidic pH + high [Cl^-^] (pH 5.7/250 mM NaCl) conditions (Table S1).

**Figure 1.**
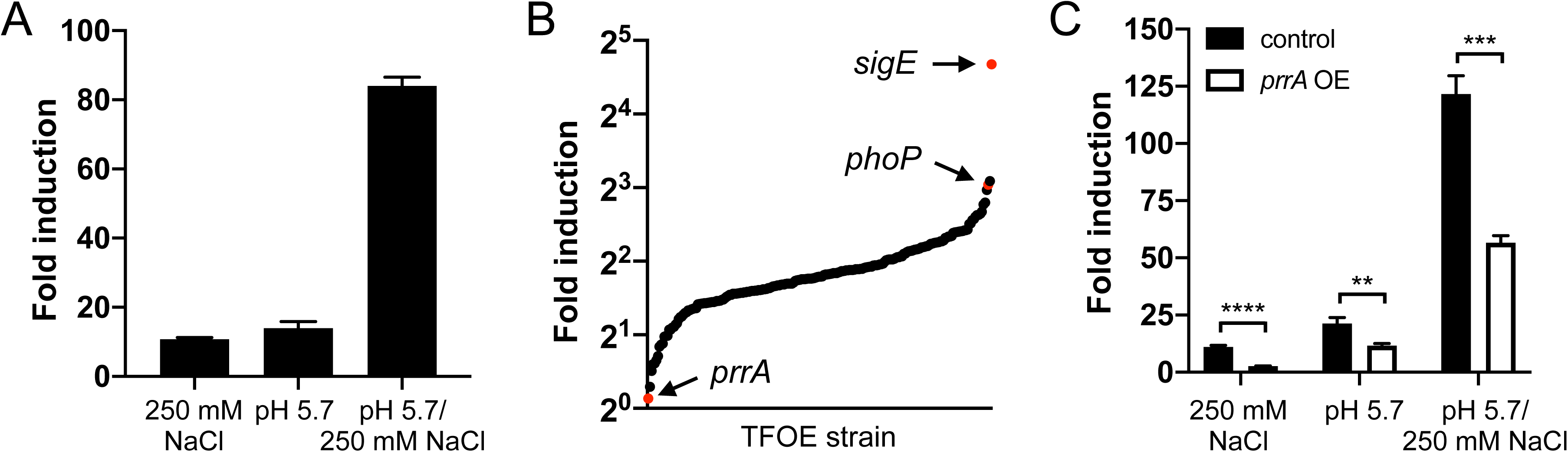
A *rv2390c’*::luciferase reporter transcription factor overexpression screen identifies PrrA as a TF that regulates Mtb response to pH and Cl^-^. (A) Mtb carrying a chromosomal *rv2390c’*::luciferase reporter was grown in 7H9, pH 7.0 ± 250 mM NaCl, or 7H9, pH 5.7 ± pH 250 mM NaCl for 9 days before light output (relative light units, RLU) and OD_600_ were measured. Fold induction compares RLU/OD_600_ in each condition to RLU/OD_600_ in the control pH 7.0 condition. Data are shown as means ± SD from three experiments. (B) A library of inducible TF overexpression plasmids (P_1_’::*TF*-FLAG-tetON) in the background of a *rv2390c’*::luciferase reporter Mtb strain was screened for their response to 250 mM NaCl. TF overexpression was induced with 200 ng/ml ATC 1 day prior to exposure to 7H9, pH 7.0 ± 250 mM NaCl media for 9 days, in the continued presence of ATC. RLU/OD_600_ was measured and fold induction calculated as in (A) for each strain. Empty vector plasmid controls were included for comparison. (C) *prrA* overexpression represses Mtb response to acidic pH and high [Cl^-^]. Mtb(P_1_’::*prrA*-FLAG- tetON, *rv2390c’*::luciferase) was grown in 7H9, pH 7.0 ± 250 mM NaCl, or 7H9, pH 5.7 ± pH 250 mM NaCl for 9 days, with ethanol (EtOH) as a carrier control (“control”) or 200 ng/ml ATC (“*prrA* OE”) added 6 days post-assay start. RLU/OD_600_ was measured at the end of the assay and fold induction calculated as in (A). Data are shown as means ± SD from three experiments. p-values were obtained with an unpaired t-test. ** p<0.01, *** p<0.001, **** p<0.0001.

On the other end of the spectrum, overexpression of *prrA* most strongly repressed induction of the *rv2390c’*::luciferase reporter in the presence of high [Cl^-^] (Fig. 1B, Table S1). While *prrA* overexpression did not result in a difference in RLU/OD_600_ values compared to the empty vector control in the acidic pH and acidic pH + high [Cl^-^] screen conditions, closer examination intriguingly indicated a decrease both in RLU and OD_600_ in the acidic pH + high [Cl^-^] condition, and an apparent increase in growth in the single acidic pH condition (Table S1). Although *prrA* is an essential gene that is part of the PrrAB two component system and has been reported to be important for Mtb replication within macrophages [35, 36], little is known about its possible role in regulating Mtb environmental response. We thus pursued further characterization of PrrA as a hit from the screen, and first validated the screen results by examining *rv2390c’*::luciferase expression in the inducible *prrA* overexpression strain (P_1_’::*prrA*-tetON, *rv2390c’*::luciferase) in uninduced (ethanol (EtOH) as a carrier control) versus induced (200 ng/ml anhydrotetracycline (ATC)) conditions. As shown in Fig. 1C, overexpression of *prrA* indeed dampened Mtb response to high [Cl^-^], and also dampened Mtb response to acidic pH alone and the dual conditions of acidic pH + high [Cl^-^]. Examination by quantitative real-time PCR (qRT-PCR) and western blot further demonstrated the expected upregulation of *prrA* transcript and protein levels upon ATC induction (Fig. S1). Together, these results suggest that PrrA acts in the regulation of Mtb response to pH and Cl^-^.

### PrrA globally regulates Mtb response to pH and Cl*^-^*

As a first follow up to the screen results with *prrA* overexpression dampening induction of the Cl^-^/pH-responsive *rv2390c’*::luciferase reporter, we pursued RNA sequencing (RNAseq) analysis to determine the global impact of PrrA on Mtb response to acidic pH and high [Cl^-^]. An initial striking observation from the RNAseq dataset was the minimal effect of *prrA* overexpression on global Mtb gene expression in the control pH 7.0 condition, with just three genes exhibiting a log_2_-fold change ≥1 (p<0.05, FDR <0.01) (Fig. 2A). In contrast, overexpression of *prrA* in the context of acidic pH + high [Cl^-^] conditions significantly impacted Mtb transcriptional response, with expression of 40 and 32 differentially expressed genes increased and decreased respectively (genes differentially expressed log_2_-fold change ≥1, p<0.05 in the EtOH set upon exposure to acidic pH/high [Cl^-^]; log_2_-fold change ≥0.25 between the ATC and EtOH sets) (Fig. 2B, Table S2). qRT-PCR analysis of several differentially expressed genes validated the results observed in the RNAseq dataset (Fig. 2C). *prrA* overexpression also impacted expression of multiple genes in the context of the single conditions of acidic pH (Figs. S2A and S2B, Table S3) or high [Cl^-^] (Figs. S2C and S2D, Table S4). Focusing on the dual condition of acidic pH and high [Cl^-^], genes involved in intermediary metabolism were among those most affected by *prrA* overexpression, with for example the induction of *prpC,* a methylcitrate synthase, *prpD,* a methylcitrate dehydratase, and *rv1405c*, a methyltransferase, all decreased (Fig. 2C, Table S2) [37, 38]. The overall trend was for *prrA* overexpression to dampen the magnitude of the gene expression differences observed, such that genes induced by acidic pH/high [Cl^-^] were less induced, while genes that were repressed by acidic pH/high [Cl^-^] were correspondingly less repressed (Fig. 2B, Table S2). There were some exceptions to this, such as with *lat*, a L-lysine aminotransferase [39], whose expression was further induced in acidic pH/high [Cl^-^] upon *prrA* overexpression (Fig. 2C, Table S2).

**Figure 2.**
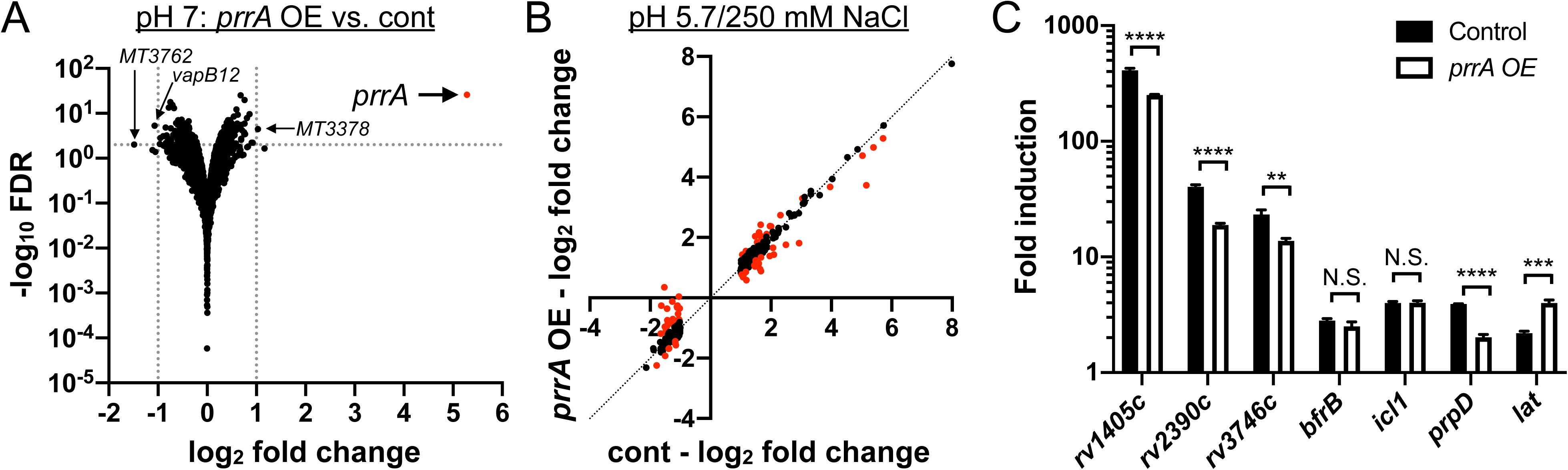
Perturbation of PrrA globally alters Mtb response to pH and Cl^-^. (A) *prrA* overexpression does not significantly alter Mtb transcriptional profile in standard pH 7.0 growth conditions. Mtb(P_1_’::*prrA*-FLAG*-*tetON, *rv2390c’*::luciferase) was grown in 7H9, pH 7.0 media and treated with EtOH or 200 ng/ml ATC for 6 hours before RNA was extracted for RNA sequencing analysis. Log_2_-fold change compares gene expression in the ATC (“*prrA* OE”) versus EtOH (“cont”) treatment sets. (B and C) *prrA* overexpression alters Mtb response to acidic pH and high [Cl^-^]. Mtb(P_1_’::*prrA*-FLAG*-*tetON, *rv2390c’*::luciferase) was grown in 7H9, pH 7.0 media and treated with EtOH or 200 ng/ml ATC for 2 hours, before exposure to 7H9, pH 7.0 or 7H9, pH 5.7 + 250 mM NaCl for 4 hours in the continued presence of EtOH or ATC as appropriate. RNA was extracted for RNA sequencing analysis (B) or qRT-PCR (C). In (B), log_2_-fold change compares gene expression in the 7H9, pH 5.7 + 250 mM NaCl condition versus the 7H9, pH 7 control condition for each of the EtOH (“cont”) or ATC (“*prrA* OE”) treatment sets. Genes marked in red had a log_2_-fold change difference ≥0.25 between the ATC and EtOH treatment sets (p<0.05, FDR<0.01 in both sets, with log_2_-fold change ≥1 in the EtOH set). In (C), fold induction compares gene expression in the pH 5.7/250 mM NaCl versus the control pH 7.0 condition for each of the EtOH (“control”) or ATC (“*prrA* OE”) treatment sets. *sigA* was used as the control gene, and data are shown as means ± SD from 3 technical replicates. p-values were obtained with an unpaired t-test. N.S. not significant, ** p<0.01, *** p<0.001, **** p<0.0001.

Together, the data reveal an environment-dependent role of PrrA in regulation of Mtb gene expression, with PrrA functioning to alter the magnitude of Mtb transcriptional response to acidic pH and high [Cl^-^].

### PrrA plays a key role in Mtb response to nitric oxide and hypoxia

Mtb experiences multiple environmental cues during host colonization, and an ability to coordinate its response to these disparate signals is critical for its adaptation within the host. Given the essentiality of PrrA and its role in modulating Mtb response to pH and Cl^-^, we thus next asked if PrrA may serve a broader function as a master TF that integrates Mtb response to multiple vital environmental cues. We first tested if PrrA plays a role in Mtb response to other ionic signals, namely iron and potassium, two ionic signals for which the bacterium has previously been shown to have a robust transcriptional response [40–43]. To test this, we incorporated a potassium-responsive *kdpF’*::GFP [43] or an iron-responsive *mbtB’*::GFP reporter plasmid into Mtb carrying a chromosomally encoded tetracycline-inducible *prrA* overexpression construct (Fig. S3A). *prrA* overexpression did not affect induction of either of these reporters (Figs. S3B and S3C), indicating that PrrA likely does not modulate Mtb response to iron or potassium. These data also support that the effects on Mtb transcriptional response to pH and Cl^-^ observed above upon *prrA* expression perturbation are specific, and not simply the result of overexpression of a gene.

Expanding beyond single ionic signals, we next tested the effect of *prrA* overexpression on Mtb response to nitric oxide (NO) and hypoxia using the NO/hypoxia-responsive *hspX’*::GFP reporter [3]. Strikingly, *prrA* overexpression significantly reduced induction of the *hspX’*::GFP reporter in response to both NO and hypoxia (Fig. 3A). To further probe the role of PrrA in Mtb response to NO, RNAseq analysis was performed on Mtb carrying the P_1_’::*prrA*-tetON plasmid, treated with EtOH (carrier control) or 200 ng/ml ATC to induce *prrA* overexpression, and exposed to 7H9, pH 7.0 media ± 100 µM DETA NONOate, an NO donor. The RNAseq data revealed that *prrA* overexpression had a marked impact on NO-mediated gene expression changes, with 80/164 genes differentially expressed log_2_-fold change ≥1 upon NO exposure demonstrating a log_2_-fold change difference ≥0.25 between the ATC and EtOH sets (Fig. 3B, Table S5). The differentially expressed genes included genes both within and outside the well-characterized DosR regulon (Figs. 3C and 3D, Table S5) [2, 4, 5]. qRT-PCR analysis showed that *dosS* induction upon NO exposure was not changed by *prrA* overexpression, although a slight decrease in *dosR* induction was noted (Fig. 3E). As observed with the acidic pH/high [Cl^-^] data above, *prrA* overexpression once again served to alter the magnitude of the transcriptional changes induced by NO; *prrA* overexpression reduced the induction of genes upregulated by NO, and dampened the extent of repression of genes downregulated by NO. Together, these data indicate the ability of PrrA to broadly repress Mtb transcriptional response to NO, with PrrA functioning to maintain proper amplitude of response of gene expression upon Mtb exposure to NO.

**Figure 3.**
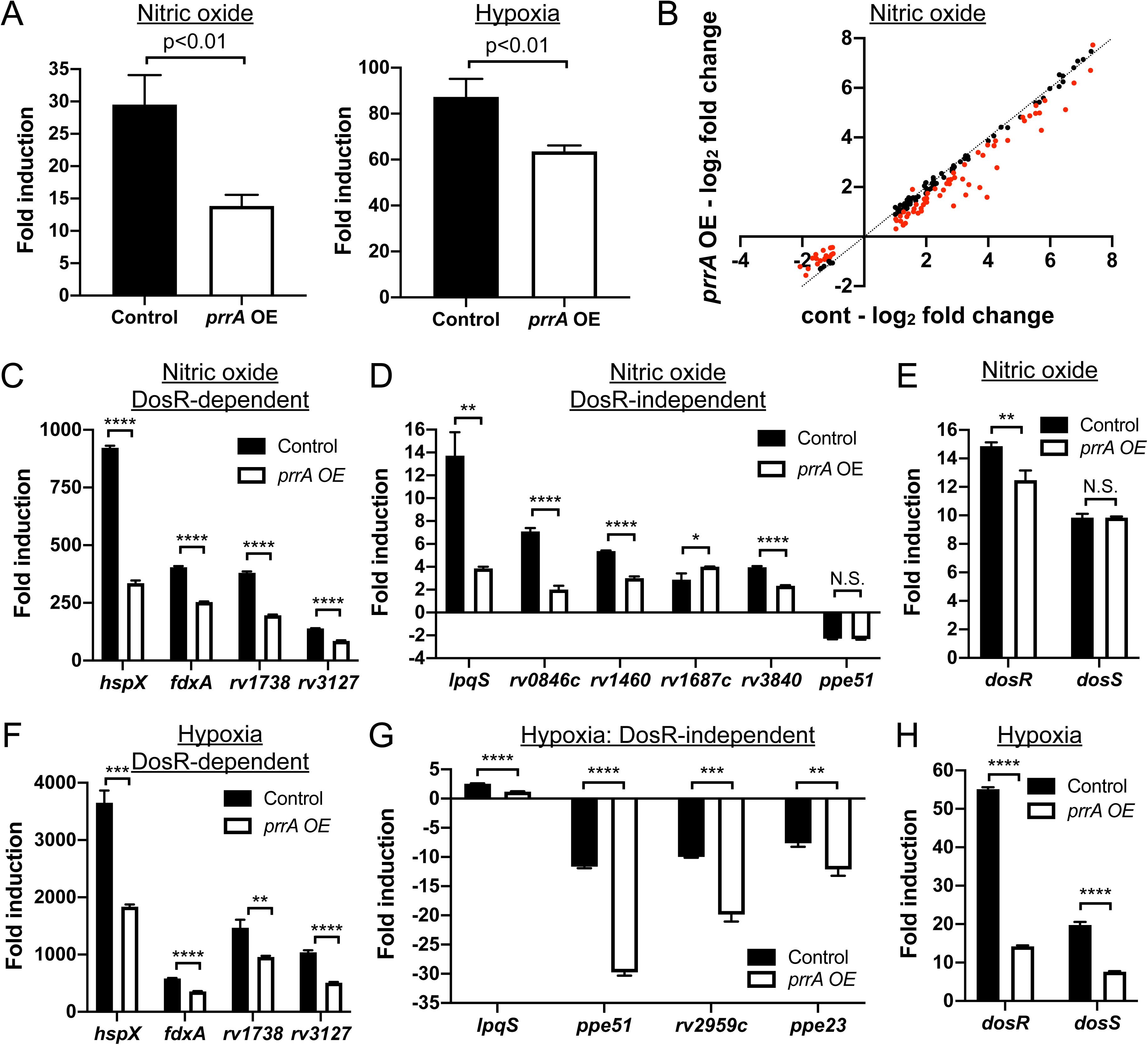
Perturbation of PrrA globally alters Mtb response to NO and hypoxia. (A) *prrA* overexpression dampens *hspX’*::GFP reporter response to NO and hypoxia. Mtb(P_606_’::*prrA*-FLAG-tetON, *hspX’::*GFP) was grown in 7H9, pH 7.0 media and treated with ETOH (“control”) or 200 ng/ml ATC (“*prrA* OE”) for 1 day before exposure to 100 µM DETA NONOate for an additional 1 day (left panel), or to 1% oxygen for 2 days (right panel). EtOH or ATC was maintained as appropriate throughout the exposure. Reporter GFP signal was analyzed by flow cytometry, and fold induction is in comparison to control conditions (no added DETA NONOate and atmospheric O_2_ respectively). Data are shown as means ± SD from 3 experiments. p-values were obtained with an unpaired t- test. (B-E) Overexpression of *prrA* globally modulates Mtb response to NO. Log-phase Mtb(P_1_’*::prrA*-FLAG*-* tetON, *rv2390c’*::luciferase) was grown in 7H9, pH 7.0 media and treated with EtOH or 200 ng/ml ATC for 2 hours, before exposure to 7H9, pH 7.0 ± 100 µM DETA NONOate for 4 hours in the continued presence of EtOH or ATC as appropriate, and samples extracted for RNA sequencing analysis (B) or qRT-PCR (C-E). In (B), log_2_-fold change compares gene expression in the 7H9, pH 7 + 100 µM DETA NONOate condition versus the 7H9, pH 7 control condition for each of the EtOH (“cont”) or ATC (“*prrA* OE”) treatment sets. Genes marked in red had a log_2_-fold change difference ≥0.25 between the ATC and EtOH treatment sets (p<0.05, FDR<0.01 in both sets, with log_2_-fold change ≥1 in the EtOH set). In (C-E), fold induction compares gene expression in the 7H9, pH 7 + 100 µM DETA-NONOate condition versus the 7H9, pH 7 control condition for each of the EtOH (“control”) or ATC (“*prrA* OE”) treatment sets. *sigA* was used as the control gene, and data are shown as means ± SD from 3 technical replicates. p-values were obtained with an unpaired t-test. N.S. not significant, * p<0.05, ** p<0.01, **** p<0.0001. (F-H) *prrA* overexpression inhibits induction of multiple hypoxia-responsive genes. qRT-PCR of Mtb(P_1_’::*prrA*-FLAG*-*tetON, *rv2390c’*::luciferase) in 7H9 pH 7.0 media treated with EtOH (“control”) or 200 ng/ ml ATC (“*prrA* OE”) for 2 hours under aerated conditions before exposure to 1% oxygen for an additional 4 hours, in the continued presence of EtOH or ATC as appropriate. Fold induction compares the 1% oxygen condition at 4 hours to the aerated 0 hour time point. *sigA* was used as the control gene, and data are shown as means ± SD from 3 technical replicates. p-values were obtained with an unpaired t-test. ** p<0.01, *** p<0.001, **** p<0.0001.

In addition to NO, overexpression of *prrA* also significantly inhibited *hspX’*::GFP induction in the presence of hypoxia (Fig. 3A). The hypoxia and NO regulons have significant overlap [4–6], and we thus anticipated that perturbation of *prrA* would also affect genes induced by hypoxia in a similar manner. Indeed, *prrA* overexpression significantly reduced hypoxia-mediated induction of both DosR-dependent and independent genes (Figs. 3F and 3G). In contrast to NO however, induction of both *dosR* and *dosS* upon hypoxia exposure was strongly reduced by *prrA* overexpression (Fig. 3H). Further investigation will be required to determine the mechanistic reasons for the differential effect of *prrA* overexpression on *dosRS* expression in hypoxia versus NO conditions.

In summary, our results establish PrrA as a master TF that regulates Mtb response not just to acidic pH and high [Cl^-^], but also to NO and hypoxia.

### PrrA, but not PrrB, is essential in Mtb

As the *prrAB* operon has previously been reported to be essential in Mtb [35], we sought to create an inducible *prrA* knockdown strain, to explore the possibility of complementary tests of the effects of *prrA* knockdown on Mtb response to environmental cues. We utilized a dual tetracycline-inducible system where ATC addition simultaneously represses expression of the gene of interest and actively degrades protein encoded by the target gene already present in the bacterium (PrrA-DUC) [44]. This construct was introduced into WT Mtb, with the native copy of *prrA* then deleted (PrrA-DUC/Δ*prrA*). Of note, we found that *prrB* was not essential, as the native *prrAB* operon could be deleted in the background of the PrrA-DUC Mtb strain (PrrA- DUC/Δ*prrAB*).

Initial characterization of the PrrA-DUC/Δ*prrA* strain showed that it grew more slowly compared to WT Mtb and expressed *prrA* at levels ∼6-fold higher, with minimal effect on *prrB* expression (Figs. 4A and 4B). PrrA protein levels were correspondingly also slightly increased in the PrrA-DUC/Δ*prrA* strain (1.36 ± 0.06 fold increase in PrrA/GroEL2 signal versus WT, Fig. 4C). Addition of ATC resulted in the expected reduction in both *prrA* transcript and PrrA protein levels, in a concentration-dependent manner (Figs. 4D and 4E). Plating for bacterial colony forming units confirmed the essentiality of *prrA* in Mtb, with survival decreasing with increasing levels of *prrA* knockdown (Fig. 4F). The sensitivity of Mtb to loss of *prrA* precluded attempts to investigate how *prrA* knockdown might affect Mtb environmental response, due to the confounding effects from the resultant bacterial death.

**Figure 4.**
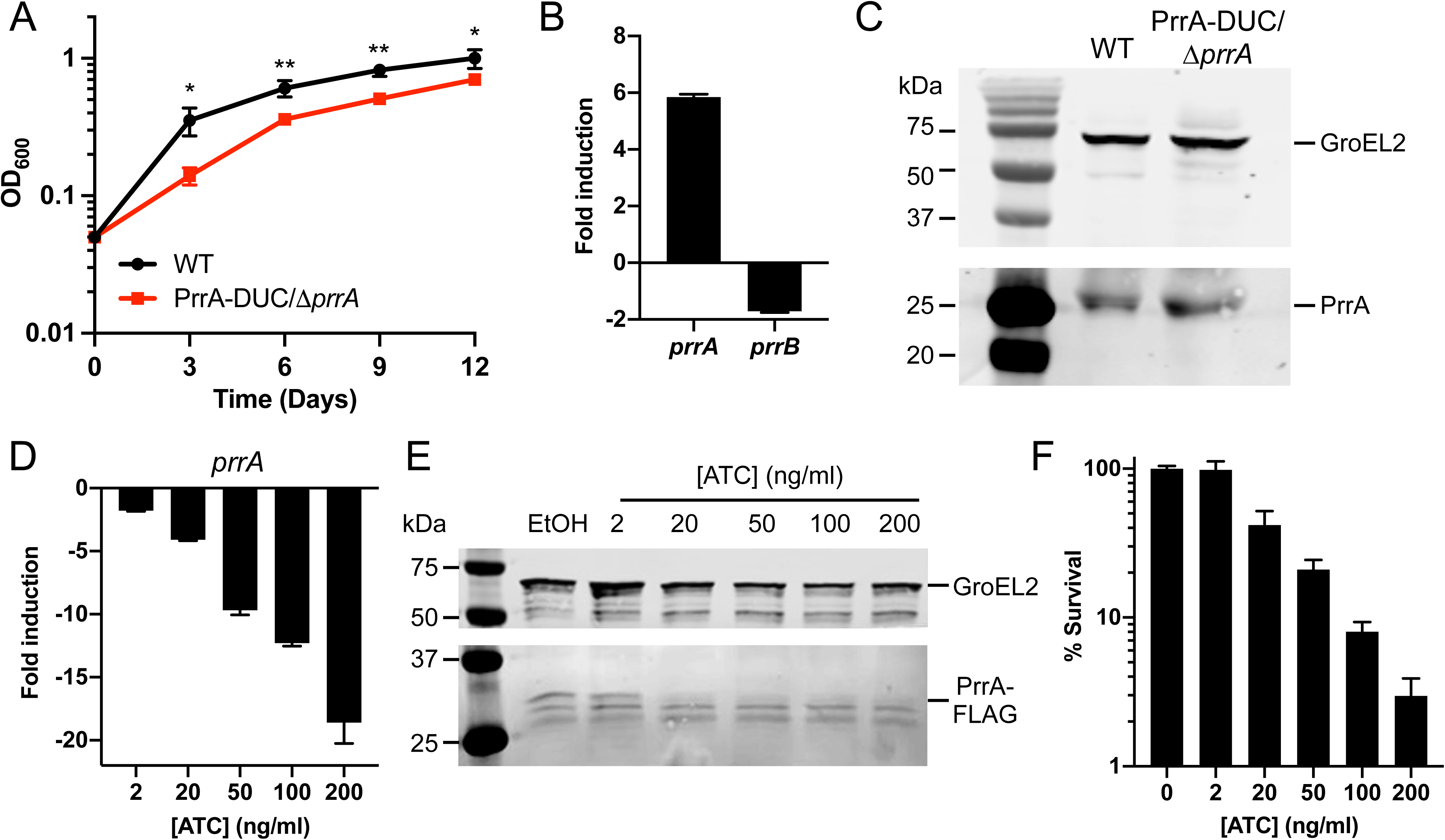
Characterization of dual inducible *prrA* gene repression/PrrA protein degradation system. (A) PrrA-DUC/Δ*prrA* Mtb grows more slowly than WT Mtb. WT or PrrA-DUC/Δ*prrA* Mtb were grown in 7H9, pH 7.0 media and growth tracked over time. Data are shown as means ± SD from three experiments. p-values were obtained with an unpaired t-test, comparing the strains at each time point. * p<0.05, ** p<0.01. (B) PrrA-DUC/Δ*prrA* Mtb has elevated *prrA* transcript levels compared to WT Mtb. qRT-PCR of WT or PrrA-DUC/Δ*prrA* Mtb grown in 7H9, pH 7.0 for four hours. Fold induction compares the PrrA-DUC/Δ*prrA* strain to WT. *sigA* was used as the control gene, and data are shown as means ± SD from three technical replicates. (C) Elevated PrrA levels is detected by western blot in PrrA-DUC/Δ*prrA* Mtb. WT or PrrA-DUC/Δ*prrA* Mtb were grown in 7H9, pH 7.0 media for nine days. Cultures were normalized to the lowest OD_600_ and lysates analyzed by western blot. Membranes were blotted with either anti-GroEL2 antibody as a loading control (top panel) or an anti-PrrA antibody (bottom panel). Blot is representative of 3 experiments. (D and E) ATC concentration-dependent repression of *prrA* expression in PrrA-DUC/Δ*prrA* Mtb. (D) shows qRT-PCR data of *prrA* expression in PrrA-DUC/Δ*prrA* Mtb grown in 7H9, pH 7.0 media and treated with EtOH or indicated concentrations of ATC for six hours. Fold induction compares each ATC-treated condition to the EtOH control. *sigA* was used as the control gene, and data are shown as means ± SD from three technical replicates. (E) shows western blot analysis of PrrA-DUC/Δ*prrA* Mtb grown in 7H9, pH 7.0 media and treated with EtOH or indicated concentrations of ATC for two days. Cultures were normalized to the lowest OD_600_ before lysate preparation. Membranes were blotted with either an anti-GroEL2 antibody as a loading control (top panel) or an anti-FLAG antibody (bottom panel). Blot is representative of 2-3 experiments. (F) Viability of the MPrrA-DUC/Δ*prrA* Mtb is affected in an ATC-concentration dependent manner. PrrA-DUC/Δ*prrA* Mtb was treated with EtOH (0 ng/ml ATC) or indicated concentrations of ATC for two days. Cultures were normalized to the lowest OD_600_ and plated for colony forming units (CFUs). CFUs observed in the control 0 ng/ml ATC concentration was set at 100% survival. Data are shown as means ± SD from three experiments.

### STPK phosphorylation plays a key role in PrrA modulation of Mtb response to pH and Cl*^-^*

Our data above reveals PrrA as an essential TF that serves as a wide-ranging modulator of Mtb response to multiple key environmental signals. Examination of *prrAB* expression showed that transcript levels of both genes were mostly unchanged in the different environmental conditions, with only hypoxia slightly upregulating *prrAB* expression (Fig. S4). In considering how PrrA activity may be differentially regulated, and in seeking a complementary method to assess the impact of *prrA* perturbation on Mtb environmental response, we thus turned to examination of its post-translational modification, in particular its phosphorylation. Of note, PrrA is phosphorylated not just by PrrB, its cognate histidine kinase (HK; D61 site), but also phosphorylated by STPKs at a second unique site (T6) [28, 45]. To investigate how phosphorylation of PrrA affects its ability to regulate Mtb response to environmental signals, we attempted to generate both HK-phosphoablative and STPK-phosphoablative PrrA variants via allelic switching at the *attL5* site [46], where the *prrA-*FLAG-DAS4 allele had been introduced in the PrrA-DUC/Δ*prrA* strain. Strikingly, we were unable to generate a HK-phosphoablative variant (PrrA-D61A). Only when the native copy of *prrA* was still present, in addition to the introduced *prrA-*FLAG-DAS4 copy, could allelic exchange with PrrA-D61A occur, indicating that phosphorylation of PrrA by a HK is essential (Fig. S5). In contrast, generation of a STPK- phosphoablative PrrA variant (PrrA-T6A) was successful, and we thus focused here on interrogating the role of STPK phosphorylation on PrrA function.

To obtain a global view of the impact of STPK phosphorylation of PrrA on PrrA regulation of Mtb response to acidic pH and high [Cl^-^], RNAseq was performed with the parental PrrA- DUC/Δ*prrA* and the PrrA-T6A-DUC/Δ*prrA* variant strain (from here on referred to as “PrrA- T6A”) exposed to pH 7.0 media (control) or to acidic pH/high [Cl^-^] media (pH 5.7/250 mM NaCl) for 4 hours. Unlike with overexpression of *prrA*, the PrrA-T6A variant impacted expression of multiple genes in standard pH 7.0 broth, with 13 genes increased in expression and 45 repressed (log_2_-fold change ≥1, p<0.05, FDR<0.01, Fig. 5A, Table S6). This suggests that the ability of PrrA to change its phosphorylation state plays a critical role in its regulation of gene expression in Mtb. Focusing on genes in the pH/Cl^-^ regulon that were differentially expressed in both WT Mtb and the parental PrrA-DUC/Δ*prrA* strain, the PrrA-T6A variant repressed 37 genes and increased the expression of 29 genes (Fig. 5B, Table S7). Of note, there were distinct differences in the effect of the PrrA-T6A variant on gene expression in response to acidic pH/high [Cl^-^] as compared to *prrA* overexpression. For example, *rv1405c* and *rv2390c* induction were not inhibited by the PrrA-T6A variant unlike with *prrA* overexpression, while expression of *bfrB,* a gene involved in iron storage and homeostasis, and *icl1,* a critical enzyme in the glyoxlate shunt pathway, were significantly down- and upregulated by the PrrA-T6A variant respectively, when *prrA* overexpression had not affected expression of these genes (Fig 5C, compare to Fig. 2C; Table S7) [47, 48]. In the case of *lat,* a gene involved in L-lysine metabolism that is upregulated in starvation and other growth inhibiting conditions [39, 49, 50], induction by acidic pH/high [Cl^-^] was significantly repressed by the PrrA-T6A variant, versus the increased expression observed upon *prrA* overexpression (Fig. 5C, compare to Fig. 2C; Table S7).

**Figure 5.**
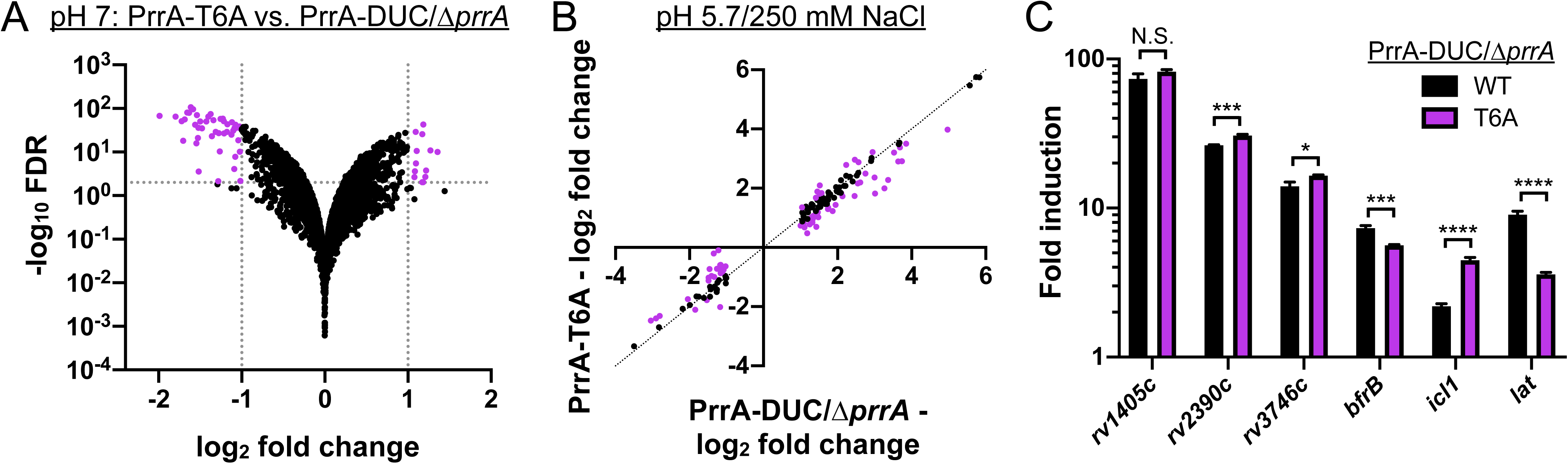
STPK phosphorylation of PrrA is important for Mtb response to pH and Cl^-^. (A) Blocking STPK phosphorylation of PrrA significantly alters Mtb transcriptional profile in standard 7H9, pH 7.0 growth conditions. PrrA-DUC/Δ*prrA* and PrrA-T6A-DUC/Δ*prrA* Mtb were grown in 7H9, pH 7.0 media for four hours before RNA was extracted for RNA sequencing analysis. Log_2_-fold change compares gene expression in the PrrA-T6A-DUC/ Δ*prrA* (“PrrA-T6A”) versus PrrA-DUC/Δ*prrA* strain (p<0.05, FDR<0.01). (B and C) A STPK phosphoablative PrrA variant alters Mtb response to acidic pH and high [Cl^-^]. PrrA-DUC/Δ*prrA* and PrrA-T6A-DUC/Δ*prrA* strains were grown in 7H9, pH 7.0 or 7H9, pH 5.7 + 250 mM NaCl for four hours, and RNA extracted for RNA sequencing analysis (B) or qRT-PCR (C). In (B), log_2_-fold change compares gene expression in the 7H9, pH 5.7 + 250 mM NaCl condition versus the 7H9, pH 7 control condition for each of the PrrA-DUC/Δ*prrA* or PrrA-T6A- DUC/Δ*prrA* strains. Genes marked in purple had a log_2_-fold change difference ≥0.25 between the PrrA-T6A-DUC/ Δ*prrA* and PrrA-DUC/Δ*prrA* strains (p<0.05, FDR<0.01 in both sets, with log_2_-fold change ≥1 in the PrrA-DUC/ Δ*prrA* set). In (C), fold induction compares the 7H9, pH 5.7 + 250 mM NaCl condition to the control 7H9, pH 7.0 condition for each strain. *sigA* was used as the control gene, and data are shown as means ± SD from 3 technical replicates. p-values were obtained with an unpaired t-test. N.S. not significant, * p<0.05, *** p<0.001, **** p<0.0001.

Together, the data reinforce a role for PrrA in regulating Mtb transcriptional response to pH and Cl^-^, and reveal the importance of STPK phosphorylation for proper function of PrrA.

### STPK phosphorylation is critical in PrrA regulation of Mtb response to NO and hypoxia

Given the significant impact of inhibition of STPK phosphorylation of PrrA on its regulation of Mtb response to pH and Cl^-^, we next sought to similarly test the role of STPK phosphorylation on PrrA control of Mtb response to NO. RNAseq was performed on the PrrA- DUC/Δ*prrA* parental strain and the PrrA-T6A variant strain exposed to pH 7.0 media (control) ± 100 µM DETA NONOate for four hours. Focusing on genes differentially expressed in both WT Mtb and the PrrA-DUC/Δ*prrA* strain upon exposure to NO, a marked modulation of gene expression by the T6A variant strain was observed, with transcript levels of 100/151 genes altered (Fig. 6A, Table S8). Strikingly, the PrrA-T6A variant dampened NO-mediated upregulation of most genes within the DosR regulon, including *dosR* and *dosS* (Figs. 6B and 6C, Table S8). The effect of the PrrA-T6A variant on DosR-independent NO-responsive genes varied, and as observed with the pH/Cl^-^ regulon, there were distinct differences in the effect of the PrrA-T6A variant on gene expression as compared to *prrA* overexpression (Fig. 6D, Table S8). For example, induction of *lpqS* and *rv0846c* by NO were both increased by PrrA-T6A, versus downregulated upon *pprA* overexpression (Fig 6D, compare to Fig. 3D; Table S8). Conversely, induction of *rv1687c* by NO was significantly reduced in the PrrA-T6A variant, versus further upregulated by *prrA* overexpression (Fig. 6D, compare to Fig. 3D; Table S8).

**Figure 6.**
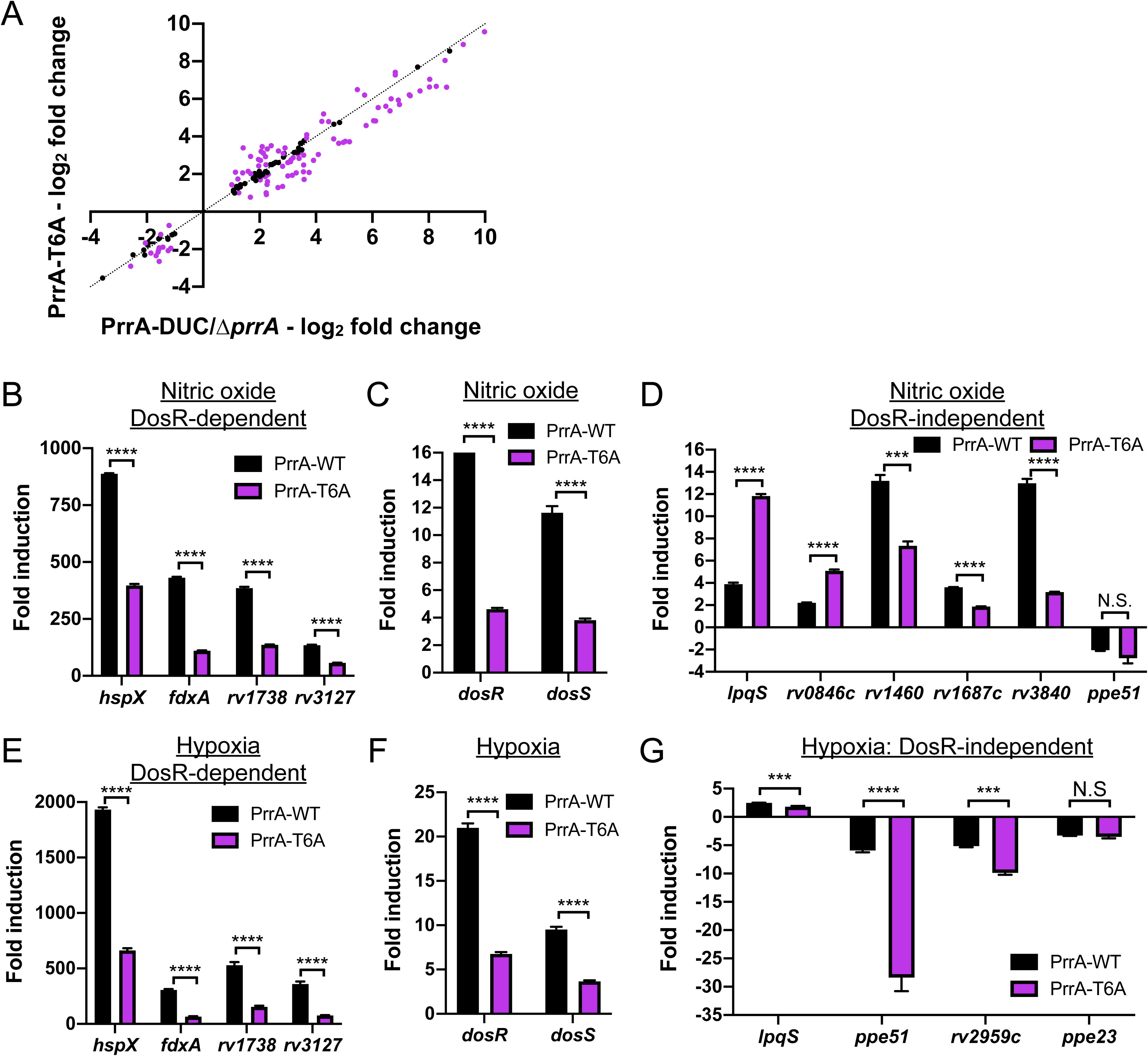
STPK phosphorylation of PrrA is critical for Mtb response to NO and hypoxia. (A-D) A STPK- phosphoablative PrrA variant strongly affects Mtb response to NO. PrrA-DUC/Δ*prrA* and PrrA-T6A-DUC/Δ*prrA* (“PrrA-T6A”) Mtb were grown in 7H9, pH 7.0 ± 100 µM DETA NONOate for four hours. RNA was extracted for RNA sequencing analysis (A) or qRT-pCR (B-D). In (A), log_2_-fold change compares gene expression in the 7H9, pH 7.0 + 100 µM DETA NONOate condition versus the 7H9, pH 7.0 control condition. Genes marked in purple had a log_2_-fold change difference ≥0.25 between the PrrA-T6A-DUC/Δ*prrA* and PrrA-DUC/Δ*prrA* strains (p<0.05, FDR<0.01 in both sets, with log_2_-fold change ≥1 in the PrrA-DUC/Δ*prrA* set). In (B-D), fold induction compares the 7H9, pH 7 + 100 µM DETA NONOate condition versus the 7H9, pH 7 control condition. *sigA* was used as the control gene, and data are shown as means ± SD from 3 technical replicates. p-values were obtained with an unpaired t-test. N.S. not significant, *** p<0.001, **** p<0.0001. (E-G) The PrrA-T6A variant strain modulates Mtb response to hypoxia. qRT-PCR of PrrA-DUC/Δ*prrA* (“PrrA-WT”) and PrrA-T6A-DUC/Δ*prrA* (“PrrA-T6A”) were grown under aerated conditions before exposure to 1% oxygen for four hours. Fold induction compares the 1% oxygen condition at 4 hours to the aerated 0 hour time point. *sigA* was used as the control gene, and data are shown as means ± SD from 3 technical replicates. p-values were obtained with an unpaired t-test. N.S. not significant, *** p<0.001, **** p<0.0001.

Parallel to the NO regulon analysis, we observed that the PrrA-T6A variant significantly inhibited hypoxia-mediated induction of multiple DosR-dependent genes, as well as *dosR* and *dosS* themselves (Figs. 6E-6F). In this case, the effect of the PrrA-T6A variant on the four DosR-independent hypoxia regulon genes generally mirrored that of *prrA* overexpression, decreasing induction of *lpqS*, and further repressing *ppe51* and *rv2959c* expression (Fig. 6G, compare to Fig. 3G). Of note, the PrrA-T6A strain had differential effects on the two DosR-independent genes whose expression are altered by both NO and hypoxia. *lpqS* induction was decreased by PrrA-T6A in the context of hypoxia exposure but increased in NO, while *ppe51* was further repressed by PrrA-T6A in the context of hypoxia exposure but unaffected in the context of NO (Figs. 6D and 6G).

Together, these data indicate that STPK phosphorylation of PrrA differentially affects PrrA activity depending on the environmental signal encountered. Importantly, there was minimal difference in the abundance of PrrA-T6A transcript or protein levels as compared to WT PrrA in the parental PrrA-DUC/Δ*prrA* strain (Fig. S6), and thus the observed phenotypes were not attributable to differences in *prrA* expression. These results reinforce the ability of PrrA to globally modulate Mtb response to pH, Cl^-^, NO, and hypoxia, and further highlight the important role of STPK phosphorylation on gene expression regulation by PrrA.

### STPK phosphorylation of PrrA is critical for Mtb entry into an adaptive state of growth arrest upon extended NO exposure

Given the overall broad inhibition of Mtb response to NO by the PrrA-T6A variant, we sought to test what the functional consequences of a failure of STPK phosphorylation of PrrA might be for the bacteria in the context of NO exposure. In particular, extended exposure to NO is known to drive Mtb into an adaptive state of growth arrest [2, 13, 51], and we thus tested the PrrA- T6A variant for its effect on Mtb entry and recovery from NO-mediated growth arrest. For this assay, the PrrA-DUC/Δ*prrA* parental strain and the PrrA-T6A variant were grown in aerated pH 7 conditions ± 100 µM DETA NONOate (four treatments across 30 hours). A first observation was that the PrrA-T6A variant grew significantly faster than its PrrA-DUC/Δ*prrA* parental strain (Fig. 7A). Strikingly, the PrrA-T6A variant also grew even upon extended NO exposure, exhibiting little growth arrest compared to the parental PrrA-DUC/Δ*prrA* strain carrying a WT allele of *prrA* (Fig. 7A). This phenotype is in accord with the significantly reduced transcriptional response of the PrrA-T6A variant to NO (Fig. 6) and demonstrates the critical role of STPK phosphorylation of PrrA in coordinating Mtb growth and NO response.

**Figure 7.**
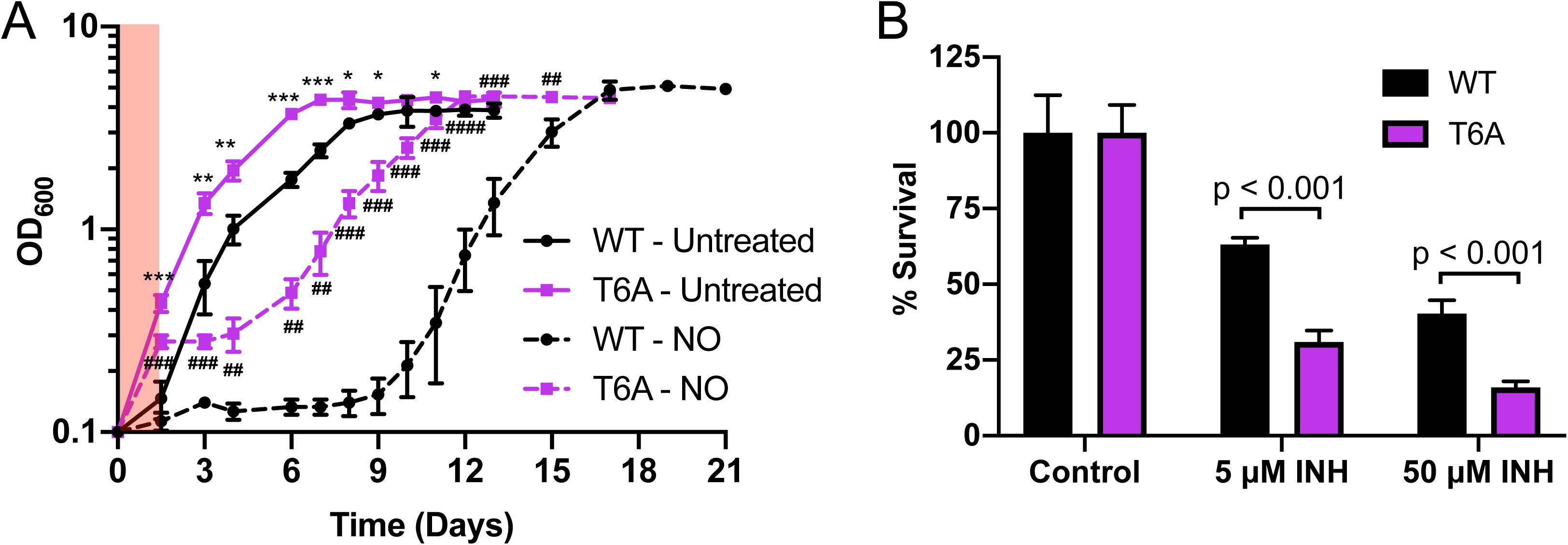
STPK phosphorylation of PrrA is critical for Mtb entry into an adaptive state of growth arrest upon extended NO exposure. (A) Blocking STPK phosphorylation of PrrA affects the ability of Mtb to enter a state of growth arrest upon extended NO exposure. PrrA-DUC/Δ*prrA* (“WT”) and PrrA-T6A-DUC/Δ*prrA* (“T6A”) were treated with 4 doses of 100 µM DETA NONOate over 30 hours (red shaded region indicates period of treatment) and growth tracked over time. Data are shown as means ± SD from 3 experiments. p-values were obtained with an unpaired t-test, comparing PrrA-T6A-DUC/Δ*prrA* to PrrA-DUC/Δ*prrA* within each condition. * symbols indicate p-values for the untreated conditions. # symbols indicate p-values for the NO conditions. * p<0.05, ** p<0.01, *** p<0.001, **** p<0.0001. (B) The PrrA-T6A variant is more sensitive to isoniazid. Log-phase PrrA-DUC/Δ*prrA* (“WT”) and PrrA-T6A-DUC/Δ*prrA* (“T6A”) Mtb were treated with isoniazid (INH) for 24 hours. Percent survival compares the number of CFUs from each INH treatment to the untreated control condition for each strain. Data are shown as means ± SD from three experiments. p-values were obtained with an unpaired t- test.

We hypothesized that the faster growth of the PrrA-T6A Mtb variant would result in increased susceptibility of the bacteria to the front-line anti-tubercular drug isoniazid (INH), which acts via inhibition of mycolic acid synthesis and has previously been shown to have increased efficacy against actively replicating Mtb [52–54]. Indeed, exposure of the PrrA-T6A variant and its parental Mtb strain to INH for 24 hours prior to plating for colony forming units (CFUs) and determining bacterial survival showed that the PrrA-T6A variant was significantly more sensitive to INH (Fig. 7B).

These results demonstrate the functional impact that STPK phosphorylation of PrrA has for Mtb, with alterations to the bacterium’s ability to respond to NO resulting in failure to adaptively adjust its replication in the presence of this growth-arresting signal. It also intriguingly suggests a role of STPK phosphorylation of PrrA in fundamental control of Mtb growth rate.

### PrrA is critical for successful host colonization by Mtb

The wide-ranging impact of PrrA on Mtb response to multiple environmental cues, and its role in enabling adaptive Mtb entry into NO-mediated growth arrest, strongly suggests that PrrA will have critical importance in the ability of Mtb to colonize its host. To directly examine this, we tested if PrrA is essential for Mtb growth within macrophages and in a murine model of Mtb infection. As expected given the essentially of *prrA*, knockdown of *prrA* significantly attenuated Mtb growth both within macrophages and mice (Figs. 8A and 8B). Similar to our observations in broth culture (Fig. 7A), preventing phosphorylation of PrrA by STPKs increased the growth rate of Mtb in macrophages (Fig. 8A). Interestingly, uptake of the PrrA-T6A Mtb variant by macrophages was significantly higher (Fig. 8A), even though the CFU/ml of the input preparation of the strain was not different from the parental PrrA-DUC/Δ*prrA* strain carrying a WT allele of *prrA* (Fig. S7). Most strikingly, the PrrA-T6A variant was significantly attenuated in its ability to colonize a whole animal host at two weeks post-infection (Fig. 8B). This indicates that proper regulation of PrrA activity, and in particular STPK phosphorylation of PrrA, is critical for Mtb host colonization *in vivo*.

**Figure 8.**
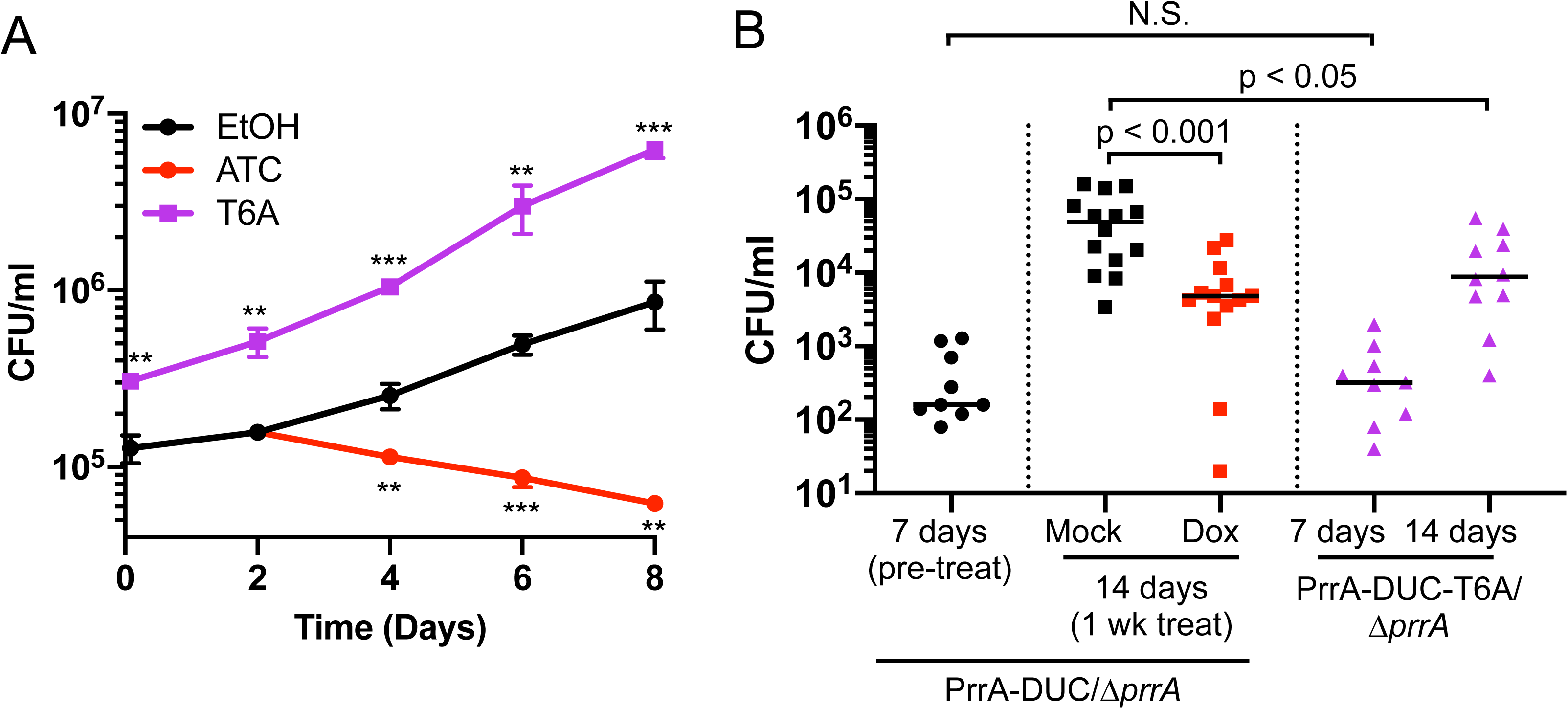
PrrA is critical for successful host colonization by Mtb. (A) *prrA* perturbation affects Mtb colonization of primary macrophages. Murine bone marrow-derived macrophages were infected with PrrA-DUC/Δ*prrA* or PrrA-T6A-DUC/Δ*prrA* (“T6A”) Mtb. EtOH or 200 ng/ml ATC was added 2 days post-infection for the PrrA-DUC/Δ*prrA* strain, with media replenished every 2 days for all strains. Data are shown as means ± SD from 3 wells. p-values were obtained with an unpaired t-test, comparing ATC-treated PrrA-DUC/Δ*prrA* or PrrA-T6A- DUC/Δ*prrA* to EtOH-treated PrrA-DUC/Δ*prrA*. ** p<0.01, *** p<0.001. (B) *prrA* perturbation inhibits Mtb colonization of mice. C3HeB/FeJ mice were infected withPrrA-DUC/Δ*prrA* or PrrA-T6A-DUC/Δ*prrA* Mtb. PrrA- DUC/Δ*prrA* Mtb infected mice were provided drinking water supplemented with 5% sucrose (“mock”) or 5% sucrose + 1 mg/ml doxycycline (“dox”) one-week post-infection for an additional week (two weeks total). Mice were sacrificed and lungs homogenized and plated for bacterial load at indicated time points. Each data point represents an individual mouse. Horizontal lines depict the median. p-values were obtained with a Mann-Whitney statistical test.

These data further emphasize the importance of proper control of bacterial growth for successful host colonization by Mtb, with a faster rate of growth *in vitro* not necessarily translating to improved outcome for the bacteria *in vivo*. Our results underscore the essential role of PrrA in Mtb infection biology and demonstrate how STPK-TCS interplay can have major impact on Mtb host colonization.

## DISCUSSION

The ability of Mtb to respond simultaneously to multiple, disparate environmental cues within its local environment is critical to its survival within a host. Here, we combined a luminescent pH/Cl^-^ transcriptional reporter strain with a tetracycline-inducible TF overexpression plasmid library to reveal the essential TF, PrrA, the response regulator (RR) of the TCS PrrAB, as a regulator that coordinates Mtb response to at least four environmental cues, namely pH, Cl^-^, NO, and hypoxia. Although the *prrAB* operon had previously been described as essential in Mtb, its functional role in Mtb biology had remained largely unknown. Early studies had indicated that *prrAB* transcript levels increased under nitrogen-limiting conditions [35], while a report utilizing a Mtb mutant with decreased *prrA* expression (transposon mutant in *prrA* promoter) suggested its importance early in macrophage infection [36]. Interestingly, *prrAB* is not essential in the non-pathogenic bacterium *Mycobacterium smegmatis*, and a study with a *M. smegmatis* Δ*prrAB* mutant reported possible roles of PrrAB in dormancy pathways [55, 56]. *M. smegmatis* has two *dosR* orthologs, and expression of *dosR2*, the ortholog located in a different genomic location from the *dosR1* and *dosS* orthologs, was decreased in the *M. smegmatis* Δ*prrAB* mutant in rich media, even in the absence of NO or hypoxia signals [55]. Unlike in Mtb, expression of several genes in the DosR regulon were additionally reported as differentially expressed in the absence of NO or hypoxia signals, with the Δ*prrAB* mutant affecting expression of a small subset of those genes [55]. These differences likely reflect fundamental differences in the role of PrrA in Mtb versus *M. smegmatis*, given the non-essentiality of PrrA in the latter.

Our transcriptional data suggests that PrrA functions as a rheostat, adjusting the amplitude of Mtb response to pH, Cl^-^, NO, and hypoxia. This contrasts with the control exerted on Mtb response to pH by PhoPR, and on a subset of the hypoxia and NO-responsive genes by DosRS(T), where deletion of these TCSs results in significantly greater inhibition of expression of the target genes more akin to an “on/off” switch [3, 5, 11]. How PrrAB, PhoPR, and DosRS(T) function may be coordinated is an intriguing question for future study. Our finding that *prrA*, but not its cognate HK *prrB*, is essential, together with our inability to generate a HK-phosphoablative PrrA variant, suggests that phosphorylation of PrrA by other HKs can occur. While TCSs are predominantly regarded as showing high specificity, with a cognate HK phosphorylating just its corresponding RR, precedence for cross-phosphorylation in TCSs exists. Examples of this include cross-phosphorylation of the HssRS and HitRS TCSs in *Bacillus anthracis* that aids in coordinating the bacterium’s response to heme and cell envelope stress [57], and PmrB phosphorylation of not just its cognate RR PmrA but also the RR QseB, part of the QseBC TCS, in the response of uropathogenic *Escherichia coli* to ferric iron [58]. Of note, a study utilizing a yeast-two hybrid approach identified the HKs DosT and PhoR as a strong and weak interactor with PrrA respectively [59]. Further studies will be required to test if PrrA can indeed be phosphorylated by these non-cognate HKs, and to reveal the cross-regulation that may exist between PrrAB, PhoPR, and DosRS(T).

Our striking findings with the STPK phosphoablative PrrA-T6A variant strongly indicates that regulation of PrrA by STPKs serve a critical role in its function, in addition to regulation by HK(s). The ability of STPKs to phosphorylate multiple targets (tens to hundreds) mark them as systems with immense capacity as masters of integrating and layering regulation. However, while STPKs (11) and TCSs (12) are both well-represented in the Mtb genome, much remains unknown about how these two vital environmental response systems may work together, with just three examples to our knowledge exploring this interplay biochemically [32, 33, 60]. The identity of the STPKs that regulate PrrA is currently unclear. A large-scale phosphoproteomic study in Mtb with follow-up assays utilizing synthetic peptides suggested two STPKs (PknD and PknF) as candidates [28], while a separate study in *M. smegmatis* using purified proteins reported a different set of three STPKs (PknG, PknJ, PknK) as being able to phosphorylate PrrA [45]. This latter study also reported using electrophoretic mobility shift assays that PrrB phosphorylation of PrrA increased DNA binding to target DNA by WT, STPK phosphoablative (T6A) and phosphomimetic (T6D) PrrA, with the strongest increase observed with the T6D PrrA variant [45]. The authors thus suggested that PrrB and STPK phosphorylation might function together in driving PrrA binding to target DNA [45]. Elucidating the STPKs that phosphorylate and regulate PrrA function will be essential for understanding the hierarchy and coordination between STPKs and HKs in Mtb environmental response and growth, and studies exploring this aspect are ongoing. More broadly, given the increasing appreciation that STPKs are not as rare in bacteria as previously thought [61], and the precedence for STPK phosphorylation of TCSs in other bacteria [62–64], we propose that future studies examining STPK-TCS interplay are likely to have important pertinence for the understanding of bacterial-host/environment interactions beyond just its relevance in Mtb.

Finally, our studies highlight the intrinsic connection between environmental signal response and Mtb growth control. Distinct from nutrient availability, Mtb growth is also strongly affected by environmental signals such as acidic pH and NO [2, 11–13], and our data with the STPK phosphoablative PrrA-T6A variant, demonstrating the inhibition of the bacterial response to NO and its failure to enter a state of growth arrest in response to NO, reinforce this aspect of Mtb biology. The intriguing ability of the STPK phosphoablative PrrA-T6A variant to grow faster than WT Mtb, combined with its attenuated *in vivo* colonization ability, lends support to the concept that Mtb is genetically “hard-wired” to slow its growth in response to certain environmental cues even in the presence of available nutrients, with rapid growth likely disadvantageous for the bacterium at least in certain stages of host infection. Further investigation of the molecular mechanisms by which STPK phosphorylation of PrrA regulates Mtb growth rate will provide vital insight into the role of STPK-TCS interplay in coordinating Mtb environmental response with its growth.

In summary, our work uncovers PrrA as an essential TF that impacts Mtb response to multiple key environmental signals, and highlights and sets the stage for future studies exploring the critical but understudied aspects of cross-regulation both among TCSs, and between STPKs and TCSs. A global understanding of how these systems interact to coordinate the bacterial response to the disparate environmental signals encountered during infection, with bacterial growth, holds significant importance in revealing master regulatory nodes that can be targeted to uncouple these two essential aspects of pathogen biology.

## MATERIALS AND METHODS

### Ethics statement

The National Institutes of Health “Guide for Care and Use of Laboratory Animals” standards were followed for all animal protocols. All animal protocols (#B2021-139) were reviewed and approved by the Institutional Animal Care and Use Committee at Tufts University, in accordance with guidelines from the Association for Assessment and Accreditation of Laboratory Animal Care, the US Department of Agriculture, and the US Public Health Service.

### Generation of Mtb strains

Strains generated in this study are listed in Table S9. Mtb cultures were maintained and propagated as described previously [11], with the following antibiotics added as appropriate: 50 µg/ml apramycin, 50 µg/ml hygromycin, 25 µg/ml kanamycin, and 100 µg/ml streptomycin. To generate the chromosomal *rv2390c’*::luciferase reporter, the *rv2390c* promoter [3] was placed upstream of the gene encoding firefly luciferase, in the integrating vector pMV306, and transformed into Mtb, with selection on 7H10 plates containing 25 µg/ml kanamycin. To construct the *mbtB’*::GFP reporter, a 638 bp region upstream of the *mbtB* start codon was PCR amplified and cloned upstream of GFPmut2 in a modified replicating plasmid (pGFP-N) as described previously for other environmental GFP reporters [3, 11]. The construct was then transformed into Mtb, with selection on 7H10 agar plates containing 50 µg/ml hygromycin. To generate P_606_’::*prrA*-FLAG- tetON, a C-terminally FLAG-tagged *prrA* driven by the P_606_ promoter was cloned via the Gateway system into the destination Gateway vector pDE43-MCK, along with a tet-ON regulator [44, 65]. To generate the dual tet-inducible *prrA*-FLAG-DAS4 knockdown system (PrrA-DUC), a C-terminally FLAG and DAS4 tagged *prrA* were cloned via the Gateway system into the destination Gateway vector pDE43-MCK (integrative plasmid). Deletion of the native copy of *prrA* or *prrB* in the background of the PrrA-DUC strain utilized homologous recombination as previously described [3]. Introduction of *prrA* point mutations was accomplished by first replacing the kanamycin marker in the pDE43-MCK vector with an apramycin marker, resulting in the plasmid pDE43-MCA. *prrA*-FLAG-DAS4 was cloned into the pDE43-MCA vector via the Gateway system, before quikchange mutagenesis (Stratagene) carried out to generate the various point mutant constructs (PrrA-T6A, PrrA-D61A). Allelic exchange at the *attL5* integration site as described in reference [46] was performed on the PrrA-DUC/Δ*prrA* Mtb strain using these new point mutant constructs, to generate PrrA-T6A-DUC/Δ*prrA* (a PrrA-D61A-DUC/Δ*prrA* strain could not be obtained as described in the Results section). All Mtb transformations were via electroporation.

### *rv2390c’*::luciferase transcription factor overexpression library construction and screen

Tetracycline-inducible transcription factor overexpression (TFOE) plasmids [23, 24] were individually introduced into the CDC1551 *rv2390c’*::luciferase reporter Mtb strain via electroporation and selected on 7H10 agar plates containing 25 µg/ml kanamycin and 50 µg/ml hygromycin. 35 µl aliquots of Mtb TFOE *rv2390c*’::luciferase strains were stored at −80°C in an arrayed format in 96-well plates, with each plate containing 23 different Mtb TFOE *rv2390c’*::luciferase strains and one empty vector control strain. To screen the library, 165 µl of 7H9, pH 7.0 media with 25 µg/ml kanamycin and 50 µg/ml hygromycin was added to thawed TFOE Mtb strains and the plates incubated for 11 days at 37°C, 5% CO_2_. TFOE *rv2390c*’::luciferase strains were then sub-cultured (1:10 dilution) into 100 µl of fresh 7H9, pH 7.0 media with appropriate antibiotics and incubated for an additional eight days at 37°C, 5% CO_2_ (to log-phase). After eight days of growth, 200 ng/ml anhydrotetracycline (ATC) was added to each well to induce overexpression of each TF for 24 hours. Afterwards, each strain was inoculated (1:10 dilution) into 100 µl of (i) 7H9, pH 7.0 (control), (ii) 7H9, pH 5.7, (iii) 7H9, pH 7, 250 mM NaCl, and (iv) 7H9, pH 5.7, 250 mM NaCl in clear-bottom, white 96-well plates (Corning Costar) and incubated for nine days at 37°C, 5% CO_2_. All media contained 50 µg/ml hygromycin and 25 µg/ml kanamycin, as well as 200 ng/ml ATC for continued TF overexpression. After nine days of incubation, luminescence was measured using the Bright-Glo luciferase assay system (Promega). Prepared luciferase substrate (75% luciferase substrate, 24.95% ultrapure water, 0.05% tween 80) was aliquoted into a v-bottom 96-well plate and 40 µl of luciferase substrate simultaneously added to each well using an Integra VIAFLO 96 liquid handler. Both light output (relative light units, RLU) and OD_600_ was measured for each well using a Biotek Synergy Neo2 multi-mode microplate reader. Fold induction was calculated by comparing RLU/OD_600_ of the samples from each environmental condition to RLU/OD_600_ of Mtb in the 7H9, pH 7.0 control condition.

To validate the results from strains of interest, the selected TFOE *rv2390c’*::luciferase Mtb strains were grown to log-phase (OD_600_ ∼0.6) in 10 ml 7H9, pH 7.0 standing, filter capped T25 flasks. Log-phase Mtb strains were then sub-cultured to an OD_600_ = 0.05 into the four media conditions described above. After six days of growth, ethanol (EtOH) as a vehicle control or 200 ng/ml ATC was added to induce overexpression of the identified TFs. Three days post-addition of EtOH or ATC, 100 µl culture aliquots in technical triplicate were transferred to a clear bottom, white 96-well plate and 40 µl of prepared luciferase substrate added/well. Light output and OD_600_ were measured, and fold induction in comparison to the 7H9, pH 7.0 control calculated in the same manner as described above.

### GFP reporter assays

Prior to the start of all GFP reporter assays, *prrA* was overexpressed for 24 hours by addition of EtOH or 200 ng/ml ATC to log-phase cultures (OD_600_ ∼0.6). These were then sub- cultured to an OD_600_ = 0.05 in the appropriate media for each reporter assay, with *prrA* overexpression maintained throughout the course of the experiment via addition of EtOH or 200 ng/ml ATC at the start of the reporter assay. At each time point, culture aliquots were taken and fixed in 4% paraformaldehyde (PFA) in phosphate buffered saline (PBS). Reporter GFP signal was determined on a BD FACSCalibur with subsequent data analysis using FlowJo (BD).

Potassium reporter assays utilizing the CDC1551(P_606_’::*prrA*-FLAG-tetON*, kdpF’*::GFP) strain were carried out in K^+^-free 7H9, pH 7.0 media supplemented with 0.05 mM KCl as described previously [43]. For iron reporter assays utilizing the CDC1551(P_606_’::*prrA*-FLAG-tetON, *mbtB’*::GFP) reporter, iron-depleted media was made essentially as previously described in Kurthkoti et al [66], except that the media was buffered to pH 7.0 with 100 mM MOPS. This iron-depleted media was supplemented with 150 µM ferric nitrate for the “control” condition, and with 100 µM 2’2’-dipyridyl (iron chelator) for the “iron-free” condition. Log-phase cultures (OD_600_ ∼0.6) in standing filter-capped T25 flasks were washed once with the iron-depleted media, then used to seed 10 ml of each of the “control” or “iron-free” media at OD_600_ = 0.05.

For NO and hypoxia reporter assays with CDC1551(P_606_’::*prrA*-FLAG-tetON, *hspX’*::GFP), Mtb cultures were first passaged three times in 4 ml filter-capped T25 flasks laid flat to aerate the cultures. Log-phase cultures (OD_600_ ∼0.6) were then sub-cultured to OD_600_ = 0.05 in 4 ml 7H9, pH 7.0 media ± 100 µM DETA NONOate (Cayman Chemicals). For hypoxia reporter assays, Mtb cultures were sub-cultured to an OD_600_ = 0.05 in 35 ml 7H9, pH 7.0 media in 125 ml filter-capped Erlenmeyer flasks (BD Biosciences), stirred with a magnetic stir bar at 45 rpm within a hypoxia chamber with adjustable O_2_ and CO_2_ controls (BioSpherix) set at 37°C, 5% CO_2_, and atmospheric O_2_. Log-phase cultures were then passaged again to an OD_600_ = 0.05 in 35 ml 7H9, pH 7.0 media in 125 ml filter-capped Erlenmeyer flasks, and incubated with a magnetic stir bar at 45 rpm within a hypoxia chamber set at 37°C, 5% CO_2_, and 1% O_2_.

### RNA sequencing and qRT-PCR analyses

For RNA sequencing (RNAseq) and qRT-PCR analyses of Mtb exposed to acidic pH and/or high [Cl^-^] conditions, log-phase Mtb (OD_600_ ∼0.6) was used to inoculate standing, filter-capped, T25 flasks containing 10 ml of (i) 7H9, pH 7.0, (ii) 7H9, pH 5.7, (iii) 7H9, pH 7.0, 250 mM NaCl, or (iv) 7H9, pH 5.7, 250 mM NaCl media at OD_600_ = 0.3. For transcriptional analysis of Mtb exposed to NO, log-phase Mtb grown in aerated conditions was used to inoculate filter-capped T75 flasks laid flat, containing 12 ml 7H9, pH 7.0 ± 100 µM DETA NONOate at OD_600_ = 0.3. Lastly, for transcriptional analysis of Mtb exposed to hypoxia, log-phase Mtb grown in aerated conditions was inoculated into 125 ml filter-capped Erlenmeyer flasks containing 35 ml 7H9, pH 7.0 media with stirring at 45 rpm and incubated at 37°C, 5% CO_2_, and 1% O_2_. For overexpression and knockdown studies, log-phase Mtb was incubated with EtOH or ATC for two hours prior to the start of the assay. Mtb samples were collected after four hours post-exposure to the indicated environmental condition, and RNA isolation was performed as previously described [1]. For RNAseq, two biological replicates per condition were used, and library preparation and sequencing carried out as previously described [43, 67]. RNAseq data was analyzed using the SPARTA program [43, 67, 68], and qRT-PCR was performed as previously described [43, 67].

### Mtb growth assays

All growth assays were performed using strains generated in the Mtb CDC1551 background. For standard growth assays, Mtb strains were grown in 10 ml 7H9, pH 7.0 media in standing, filter-capped T25 flasks with the appropriate antibiotics as previously described [3]. Log-phase Mtb strains (OD_600_ ∼0.6) were used to inoculate cultures at a starting OD_600_ = 0.05, and growth was tracked over time via OD_600_ measurement.

For isoniazid (INH) sensitivity assays, Mtb strains were grown in 10 ml 7H9, pH 7.0 media in standing, filter capped T25 flasks for 6 days at a starting OD_600_ = 0.05. After 6 days of growth, 198 µl of Mtb culture were transferred per well of a 96-well plate, and incubated in the presence of 5 and 50 µM INH for 24 hours at 37°C, 5% CO_2_. For determination of colony forming units (CFUs), strains were serially diluted in PBS + 0.05% tween 80 and plated for CFUs on 7H10 agar plates containing 100 µg/ml cycloheximide.

For NO growth arrest assays, log-phase Mtb cultures (OD_600_ ∼0.6) were passaged three times in filter-capped T75 flasks that were laid flat, in 12 ml 7H9, pH 7.0 media. After the third passage, Mtb strains were sub-cultured to OD_600_ = 0.1 in 12 ml 7H9, pH 7.0 media, in filter capped T75 flasks laid flat. Mtb was then treated with 100 µM DETA NONOate (Cayman Chemicals) 4 times across 30 hours. Growth was tracked over time via OD_600_ measurement.

### PrrA protein purification

The sequence encoding C-terminally FLAG-tagged *prrA* was cloned into the isopropyl-β-D-1- thiogalactopyranoside (IPTG)-inducible pET-23a vector and transformed into *Escherichia coli* BL21(DE3) for recombinant expression and purification of *prrA*-FLAG. For expression, 2 ml of overnight *E. coli* cultures from frozen stock grown in 5 ml LB + 50 µg/ml carbenicillin at 37°C was used to inoculate 1 L of 2x YT media + 50 µg/ml carbenicillin, and grown with shaking at 150 rpm at 37°C until the culture reached an OD_600_ of ∼0.7. To induce production of PrrA-FLAG, 1 mM IPTG was added and the cultures grown for an additional 4 hours at 37°C, 150 rpm. Afterwards, supernatants were removed and pellets were stored at −80°C prior to further processing.

Purification of PrrA-FLAG protein was based on a previously described FLAG-tagged protein immunoprecipitation protocol [69]. Thawed pellets were resuspended in 35 ml FLAG immunoprecipitation buffer (FIB) (50 mM Tris, pH 7.5, 150 mM NaCl, and protease inhibitors (Pierce protease inhibitor tablet)). Resuspended cells were lysed by sonication (3 x 45 seconds at 40% amplitude, five minutes of rest on ice between each sonication), the insoluble fraction pelleted by centrifugation at 4,000 rpm for 30 minutes at 4°C, and the cleared whole cell lysate removed into a new 50 ml conical tube. Cleared whole cell lysate was incubated with anti-FLAG M2 magnetic beads (Sigma #M8223-5ML) at room temperature for one hour, with shaking at 50 rpm. The anti-FLAG M2 magnetic beads were separated from the cleared whole cell lysate with a magnetic tube holder and beads were washed in 1 ml FIB buffer three times. After washing, anti-FLAG M2 magnetic beads were incubated with 100 µg/ml 3x FLAG peptide (Sigma, #F4799- 25MG) in FIB buffer + 10% glycerol for 30 minutes at room temperature. PrrA-FLAG protein was collected by separating the anti-FLAG M2 magnetic beads with a magnetic tube holder, and separated PrrA-FLAG protein was aliquoted into a new microcentrifuge tube. PrrA-FLAG purification was confirmed by running samples on a 12% SDS-PAGE gel, visualized by Coomassie Brilliant Blue staining, and imaged using the 700 nm channel of an Odyssey CLx imaging system. A Bradford assay was performed to quantify protein concentration.

### Western blot analysis

For detection of native PrrA in WT Mtb or basal levels of PrrA in the PrrA-DUC/Δ*prrA* and PrrA-T6A-DUC/Δ*prrA* strains, log-phase Mtb strains (OD_600_ ∼0.6) were sub-cultured to an OD_600_ = 0.05 in 10 ml 7H9, pH 7.0 media with appropriate antibiotics in standing, filter-capped T25 flasks and grown for nine days at 37°C, 5% CO_2_ before lysate collection. For measurement of PrrA overproduction in the CDC1551(P_1_’::*prrA*-FLAG-tetON, *rv2390c’*::luciferase) strain, log-phase Mtb (OD_600_ ∼0.6) was sub-cultured to an OD_600_ = 0.05 in 10 ml 7H9, pH 7.0 media with appropriate antibiotics and grown for six days at 37°C, 5% CO_2_ before induction with 200 ng/ml ATC or EtOH as a vehicle control for an additional 3 days prior to lysate collection. Lastly, for measurement of PrrA degradation with the PrrA-DUC/Δ*prrA* Mtb strain, cultures were grown for six days before addition of EtOH or varying ATC concentrations for an additional two days prior to lysate collection.

For collection of cell lysates in all cases, Mtb cultures were normalized to the lowest OD_600_ and washed twice with 1 ml wash buffer (10 mM Tris, pH 8.0, 0.05% tween 80). Pellets were stored at −80°C prior to further processing. Thawed pellets were resuspended in 900 µl lysis buffer (50 mM Tris-HCl, 1% SDS, 5 mg/ml lysozyme, 5 mM EDTA, and protease inhibitors (Pierce protease inhibitor tablet)). Resuspended cells were transferred to 2 ml tubes containing 300 µl 0.1 mm silica beads and cells were lysed via bead beating (2 x 1 min, 6 m/s) with a MP Biomedicals Fastprep-24 system. Whole cell lysates were then transferred to new 1.5 ml microcentrifuge tubes and boiled at 100°C for 15 minutes. After boiling, an appropriate amount of 4X loading buffer (LI- COR Biosciences) with β-mercaptoethanol was added to whole cell lysates followed by an additional 15 minutes of boiling at 100°C.

For western blot analysis, samples were resolved on a 12% SDS-PAGE gel and transferred to Millipore Immobilon-FL PVDF membranes. Membranes were blocked with LI-COR Odyssey buffer for 30 minutes at room temperatures and incubated with either mouse anti-Mtb GroEL2 (Clone IT-70 (DCA4), BEI Resources #NR-13657) at 1:2500, rabbit anti-PrrA (generous gift from Timothy Donohue, [70]) at 1:10,000, or mouse anti-FLAG (Sigma, #F3165-MG) at 1:1000 overnight at room temperature. Membranes were then washed 3 x 5 minutes with PBS + 0.1% tween 20 and incubated with goat anti-mouse IRDye 680CW at 1:5000 or goat anti-rabbit IRDye 680CW at 1:5000 (LI-COR) for one hour at room temperature, as appropriate. Blots were washed 3 x 5 minutes with PBS 0.1% tween 20, then rinsed with PBS, before visualization with a LI-COR Odyssey CLx imaging system. Image Studio software (LI-COR Biosciences) was used to quantify the signal intensity of protein bands.

### Macrophage culture and infections

C57BL/6J mice (The Jackson Laboratory, Bar Harbor, ME) were used for isolation of bone marrow-derived macrophages (BMDMs). BMDMs were maintained in DMEM media containing 10% FBS, 15% L-cell conditioned media, 2 mM L-glutamine, 1 mM sodium pyruvate, and penicillin/streptomycin when appropriate, in a 37°C, 5% CO_2_ incubator. BMDM infections with Mtb and subsequent enumeration of bacterial load in the macrophages were carried out as previously described [43]. To test how *prrA* knockdown affected Mtb colonization of BMDMs, 200 ng/ml ATC was added to the culture media two days post-infection and ATC was replenished every two days along with fresh culture media.

### Mouse Mtb infections

C3HeB/FeJ mice (The Jackson Laboratory) were intranasally infected with 10^3^ CFUs of Mtb (35 µl volume) while under light anesthesia with 2% isoflurane [3, 8, 71]. For infections to test how *prrA* knockdown affected host colonization, mice were given water supplemented with 5% sucrose ± 1 mg/ml doxycycline beginning one week after infection for one additional week [67, 72]. Water was replaced twice during the week of treatment. After mice were sacrificed with CO_2_, the left lobe and accessory right lobe were homogenized in PBS + 0.05% tween 80, while the remaining right lobes were fixed with 4% paraformaldehyde. Homogenates were serially diluted and plated onto 7H10 agar plates containing 100 µg/ml cycloheximide for CFU quantification.

### Accession numbers

RNA sequencing data have been deposited in the NCBI GEO database (GSE198997-GSE199000).

## ACKNOWLEDGEMENTS

We thank Yuzo Kevorkian and Richard Lavin for assistance with the mice infection studies and Natalia Quirk for initial work on PrrA purification. We thank David Sherman (University of Washington, Seattle, WA) for generously providing the tetracycline-inducible transcription factor overexpression plasmids, Dirk Schnappinger (Weill Cornell Medical School, New York, NY) for the dual tetracycline-inducible gene repression/protein degradation system, and Timothy Donohue (University of Wisconsin-Madison, Madison, WI) for providing the rabbit anti-PrrA antibody. We thank Albert Tai for assistance with RNA sequencing analysis. This work was supported by grants from the National Institutes of Health (R01 AI143768) and the American Lung Association (IA- 823328) to ST. DG was supported by a Ruth L. Kirschstein National Research Service Award from the National Institutes of Health (F31 AI161032) and by training grant T32 GM007310 to Andrew Camilli from the National Institutes of Health. The funders had no role in study design, data collection and analysis, decision to publish, or preparation of the manuscript.

## SUPPLEMENTAL TABLES

**Table S1. Results of CDC1551(*rv2390c’*::luciferase) transcription factor overexpression pH/Cl^-^ screen.**

**Table S2. Comparison of effect of *prrA* overexpression on genes differentially expressed log_2_- fold change ≥1 in the EtOH treatment set, after 4 hours exposure to pH 5.7/250 mM NaCl.**

**Table S3. Comparison of effect of *prrA* overexpression on genes differentially expressed log*_2_*- fold change ≥1 in the EtOH treatment set, after 4 hours exposure to pH 5.7.**

**Table S4. Comparison of effect of *prrA* overexpression on genes differentially expressed log_2_- fold change ≥1 in the EtOH treatment set, after 4 hours exposure to 250 mM NaCl.**

**Table S5. Comparison of effect of *prrA* overexpression on genes differentially expressed log*_2_*- fold change ≥1 in the EtOH treatment set, after 4 hours exposure to 100 µM DETA NONOate.**

**Table S6. Differentially expressed genes (log_2_-fold change ≥1, p<0.05) in PrrA-T6A- DUC/ΔprrA (“T6A”) versus PrrA-DUC/ΔprrA (“WT”) in 7H9, pH 7.0.**

**Table S7. Comparison of effect of PrrA-T6A-DUC/ΔprrA (“T6A”) versus PrrA-DUC/Δ*prrA* on genes differentially expressed ≥1 log*_2_*-fold change in the WT and PrrA-DUC/Δ*prrA* sets, after 4 hours exposure to pH 5.7/250 mM NaCl.**

**Table S8. Comparison of effect of PrrA-T6A-DUC/ΔprrA (“T6A”) versus PrrA-DUC/*ΔprrA* on genes differentially expressed ≥1 log_2_-fold change in the WT and PrrA-DUC/*ΔprrA* sets, after 4 hours exposure to 100 µM DETA NONOate.**

**Table S9. Strains generated in this study.**

## Notes

### Competing Interest Statement

The authors have declared no competing interest.

