## Supplemental Figures for "PrrA modulates *Mycobacterium tuberculosis* response to multiple environmental cues and is critically regulated by serine/threonine protein kinases"

### Figure S1

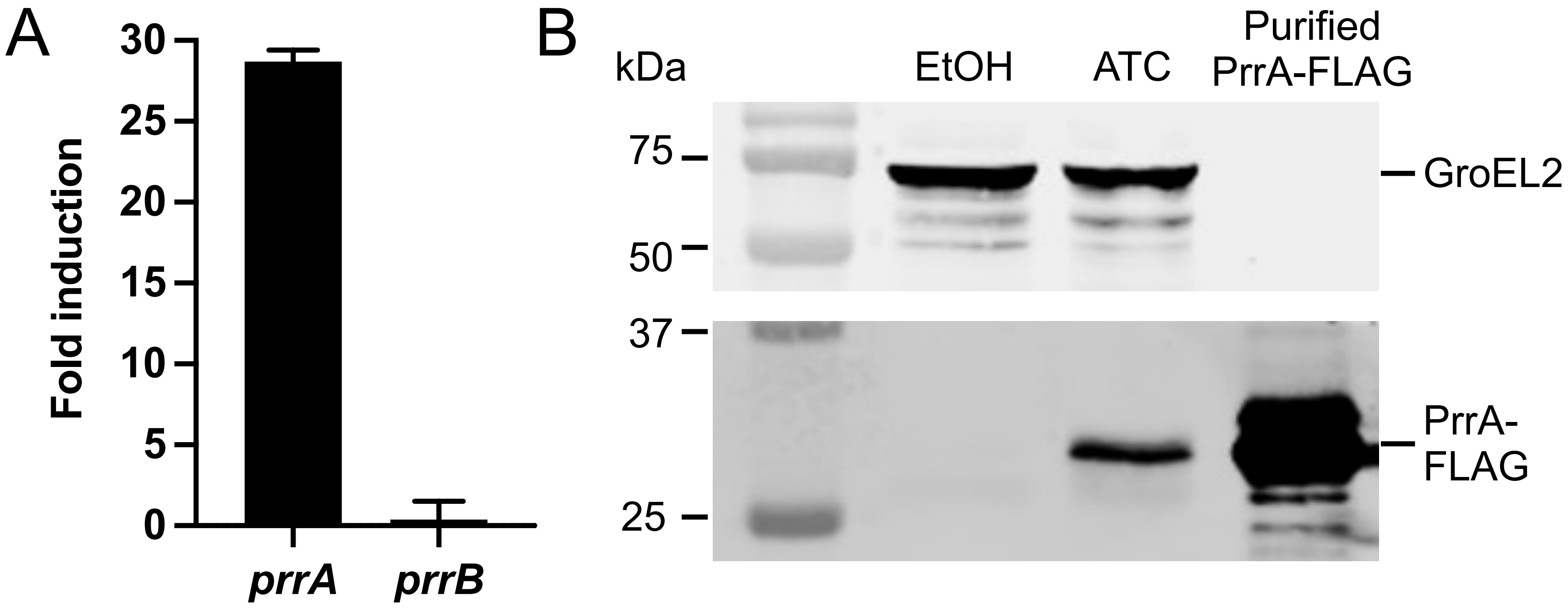


**Figure S1. Inducible *prrA* overexpression system in Mtb.** (A) qRT-PCR of Mtb($P_1$'*::prrA*-FLAG-tetON, *rv2390c'*::luciferase) treated with ethanol (EtOH) or 200 ng/ml ATC for 6 hours in 7H9, pH 7.0 media. Fold induction compares the ATC to the EtOH treatment. *sigA* was used as the control gene, and data are shown as means ± SD from 3 technical replicates. (B) Mtb($P_1$'*::prrA*-FLAG-tetON, *rv2390c'*::luciferase) was exposed to EtOH or 200 ng/ml ATC for 3 days. Cultures were then normalized to the lowest $OD_{600}$ and lysates analyzed by western blot. Membranes were blotted with either an anti-GroEL2 antibody as a loading control (top panel) or an anti-FLAG antibody (bottom panel). Purified recombinant PrrA-FLAG protein was used as a positive control. Blot is representative of 3 experiments.

### Figure S2

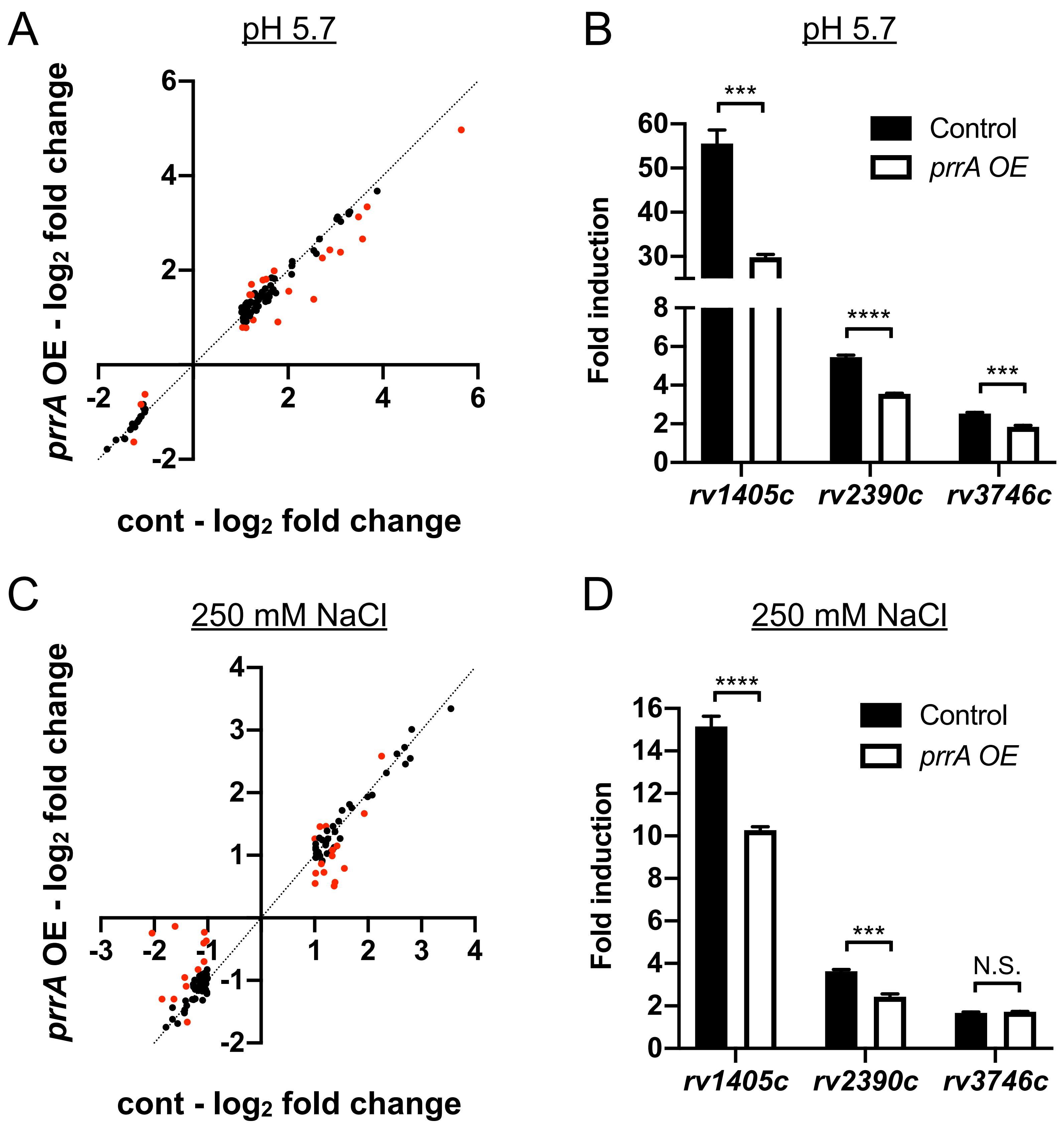


**Figure S2. Perturbation of PrrA globally alters Mtb response to single environmental cues of pH or Cl-.** (A and B) *prrA* overexpression alters Mtb response to acidic pH. Mtb($P_1$'::*prrA*-FLAG-tetON, *rv2390c'*::luciferase) was grown in 7H9, pH 7.0 media and treated with EtOH or 200 ng/ml ATC for 2 hours, before exposure to 7H9, pH 7.0 or 7H9, pH 5.7 for four hours, in the continued presence of EtOH or ATC as appropriate. RNA was extracted for RNA sequencing (A) or qRT-PCR (B) analysis. In (A), $\log_2$-fold change compares gene expression in the 7H9, pH 5.7 condition versus the 7H9, pH 7 control condition for each of the EtOH ("cont") or ATC ("*prrA* OE") treatment sets. Genes marked in red had a $\log_2$-fold change difference $\geq 0.25$ between the ATC and EtOH treatment sets ($p<0.05$, FDR<0.01 in both sets, with $\log_2$-fold change $\geq 1$ in the EtOH set). In (B), fold induction compares gene expression in the pH 5.7 versus the control pH 7.0 condition for each of the EtOH ("control") or ATC ("*prrA* OE") treatment sets. *sigA* was used as the control gene, and data are shown as means ± SD from 3 technical replicates. p-values were obtained with an unpaired t-test. *** $p<0.001$, **** $p<0.0001$. (C and D) *prrA* overexpression alters Mtb response to high [Cl-]. Mtb($P_1$'::*prrA*-FLAG-tetON, *rv2390c'*::luciferase) was grown in 7H9, pH 7.0 media and treated with EtOH or 200 ng/ml ATC for 2 hours, before exposure to 7H9, pH 7.0 ± 250 mM NaCl for four hours, in the continued presence of EtOH or ATC as appropriate. RNA was extracted for RNA sequencing (C) or qRT-PCR (D) analysis. In (C), $\log_2$-fold change compares genes expression in the 7H9, pH 7.0 + 250 mM NaCl condition versus the 7H9, pH 7.0 control condition for each EtOH ("cont") or ATC ("*prrA* OE") treatment sets. Genes are marked in red as in (A). In (D), fold induction compares gene expression in the 250 mM NaCl versus the control pH 7.0 condition for each of the EtOH ("control") or ATC ("*prrA* OE") treatment sets. Data are shown as in (B). p-values were obtained with an unpaired t-test. N.S. not significant, *** $p<0.001$, **** $p<0.0001$.

### Figure S3

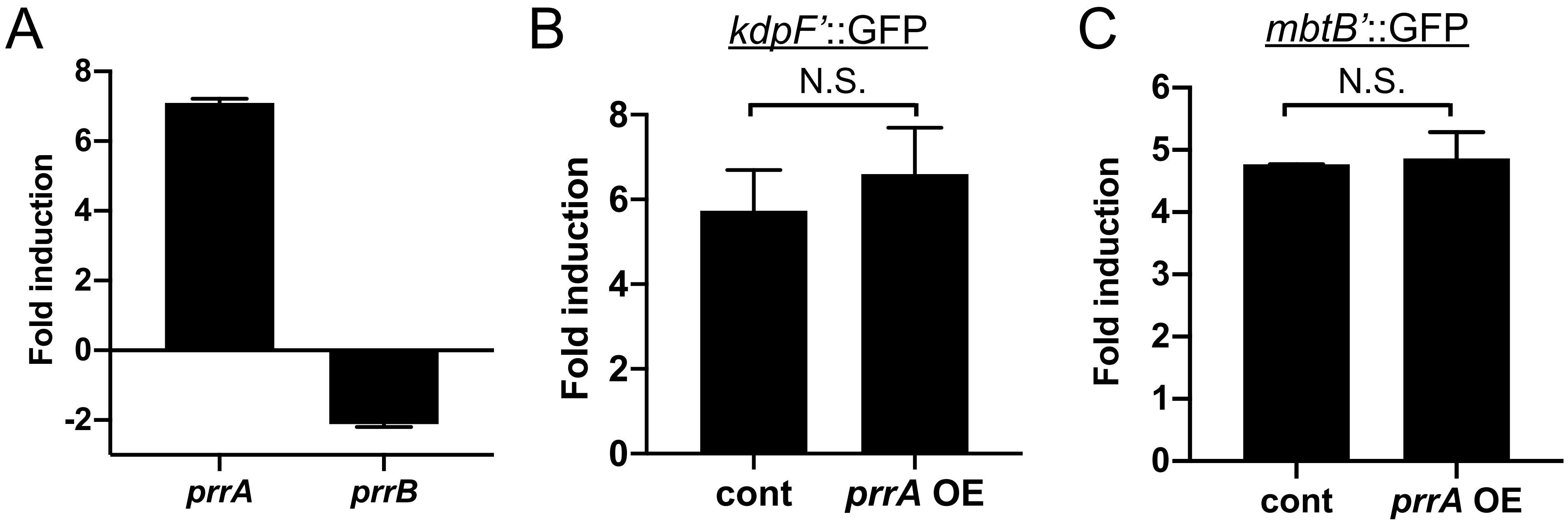


**Figure S3. *prrA* overexpression does not affect induction of potassium or iron-responsive Mtb reporters.** (A) *prrA* is inducibly overexpressed in a Mtb strain carrying a chromosomal copy of an ATC-inducible *prrA*-FLAG construct. qRT-PCR of Mtb($P_{606}$'::*prrA*-FLAG-tetON) treated with EtOH or 200 ng/ml ATC for 6 hours in 7H9, pH 7.0 media. Fold induction compares the ATC-treated condition to the EtOH control. *sigA* was used as the control gene, and data are shown as means ± SD from three technical replicates. (B) *prrA* overexpression does not affect Mtb response to low [$K^+$]. Mtb($P_{606}$'::*prrA*-FLAG-tetON, *kdpF'*::GFP) was treated with EtOH or 200 ng/ml ATC for 24 hours before exposure to 7H9, pH 7.0 or $K^+$-free 7H9, pH 7.0 media for 9 days. EtOH or ATC was maintained as appropriate throughout the exposure. Reporter GFP fluorescence was measured by flow cytometry, and fold induction is in comparison to the control 7H9, pH 7.0 condition. Data are shown as means ± SD from three experiments. p-value was obtained with an unpaired t-test. N.S. not significant. (C) *prrA* overexpression does not affect Mtb response to low iron. Mtb($P_{606}$'::*prrA*-FLAG-tetON, *mbtB'*::GFP) was treated with EtOH or 200 ng/ ml ATC for 24 hours before exposure to iron-depleted media + 150 µM $Fe(NO_3)_3$ (control) or iron-depleted media + 100 µM 2,2'-dipyridyl (iron chelator) for 9 days. EtOH or ATC was maintained as appropriate throughout the exposure. Reporter GFP fluorescence was analyzed and data presented as in (B). p-value was obtained with an unpaired t-test. N.S. not significant.

### Figure S4

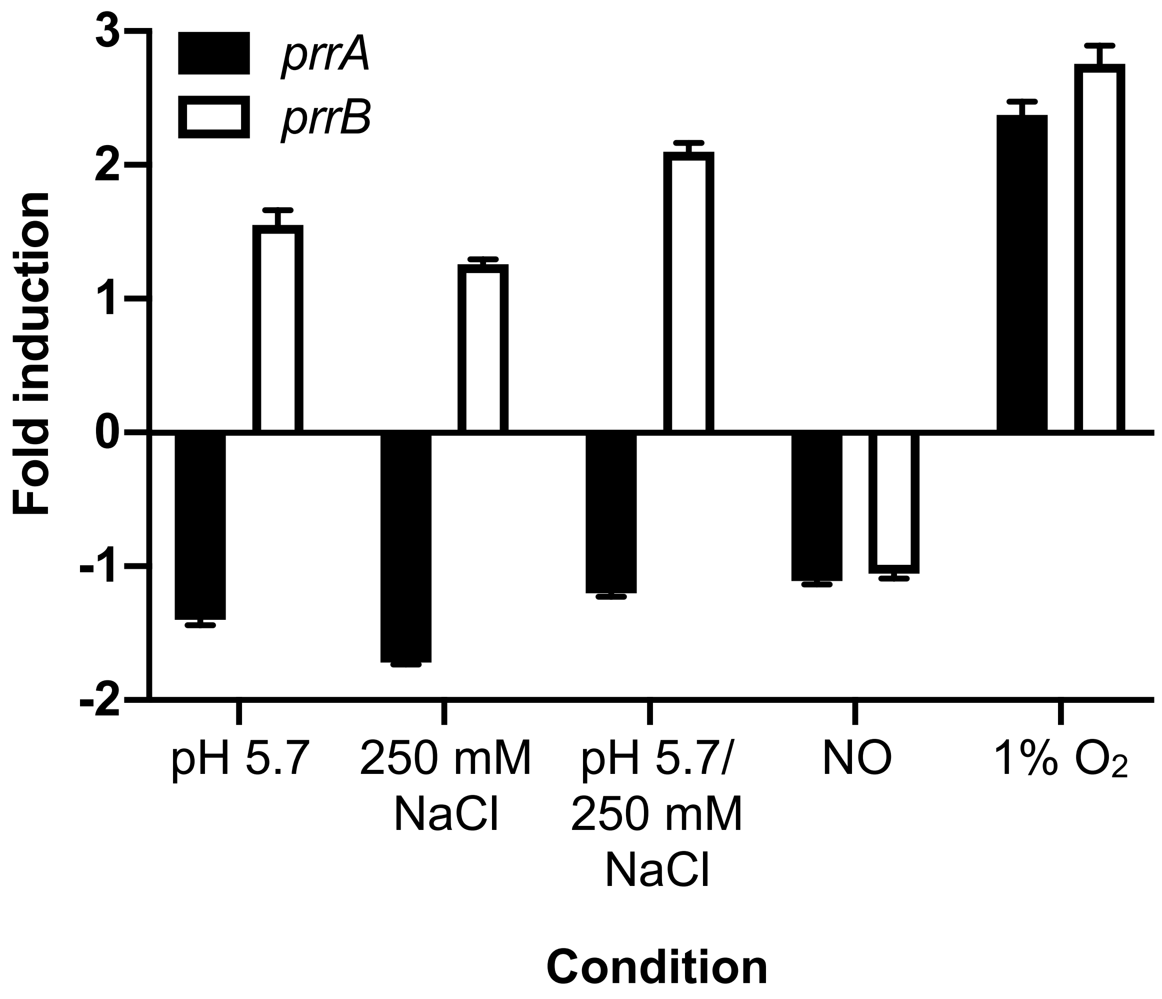


**Figure S4. *prrAB* expression is largely unchanged under various environmental conditions.** WT Mtb was exposed to each indicated environmental condition or 7H9, pH 7.0 (control) for 4 hours. qRT-PCR data is shown, with fold induction comparing each environmental condition to the 7H9, pH 7.0 control condition except for the "1% oxygen" condition, where fold induction is in comparison to the aerated 7H9, pH 7.0 condition at the 0 hour time point. *sigA* was used as the control gene, and data are shown as means ± SD from three technical replicates.

### Figure S5

PrrA-DUC | PrrA-DUC/Δ*prrA*

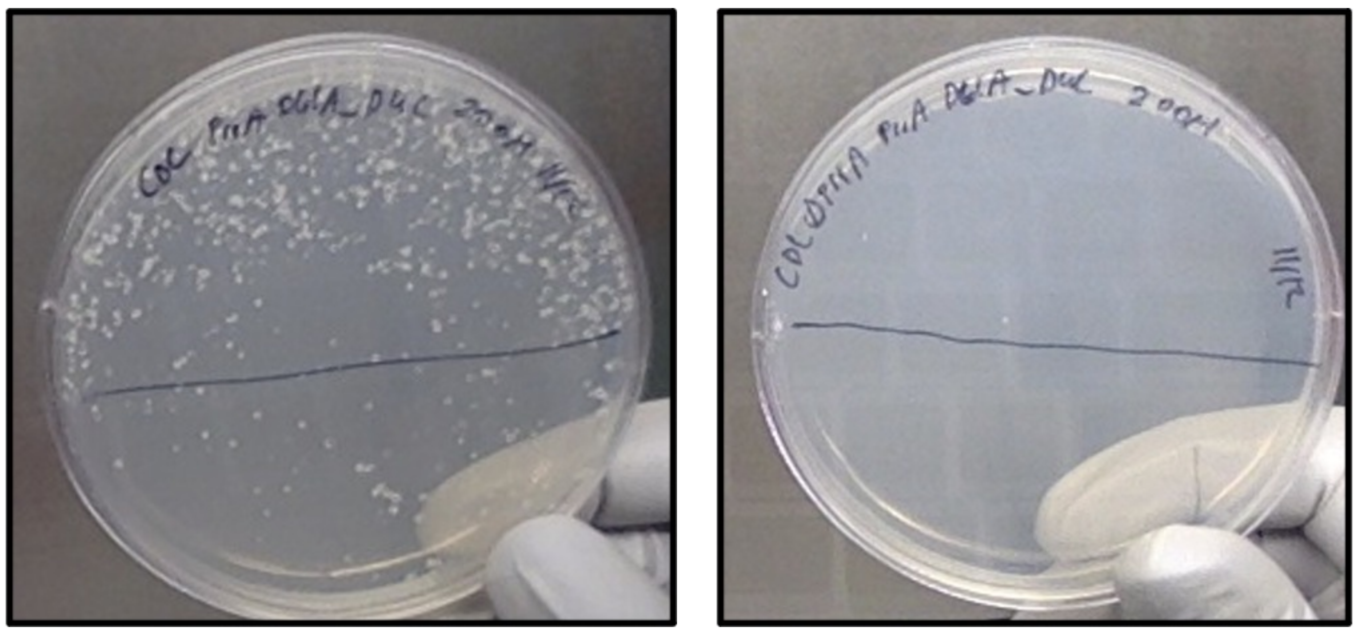

**Figure S5. Histidine kinase phosphorylation of PrrA is essential in Mtb.** Allelic exchange of Mtb strains carrying two copies of *prrA* ("PrrA-DUC") or one copy of *prrA* ("PrrA-DUC/Δ*prrA*") was attempted, to replace the WT PrrA-DUC copy with a copy containing the PrrA-D61A variant. Only when the native copy of *prrA* was present were colonies obtained (left panel).

### Figure S6

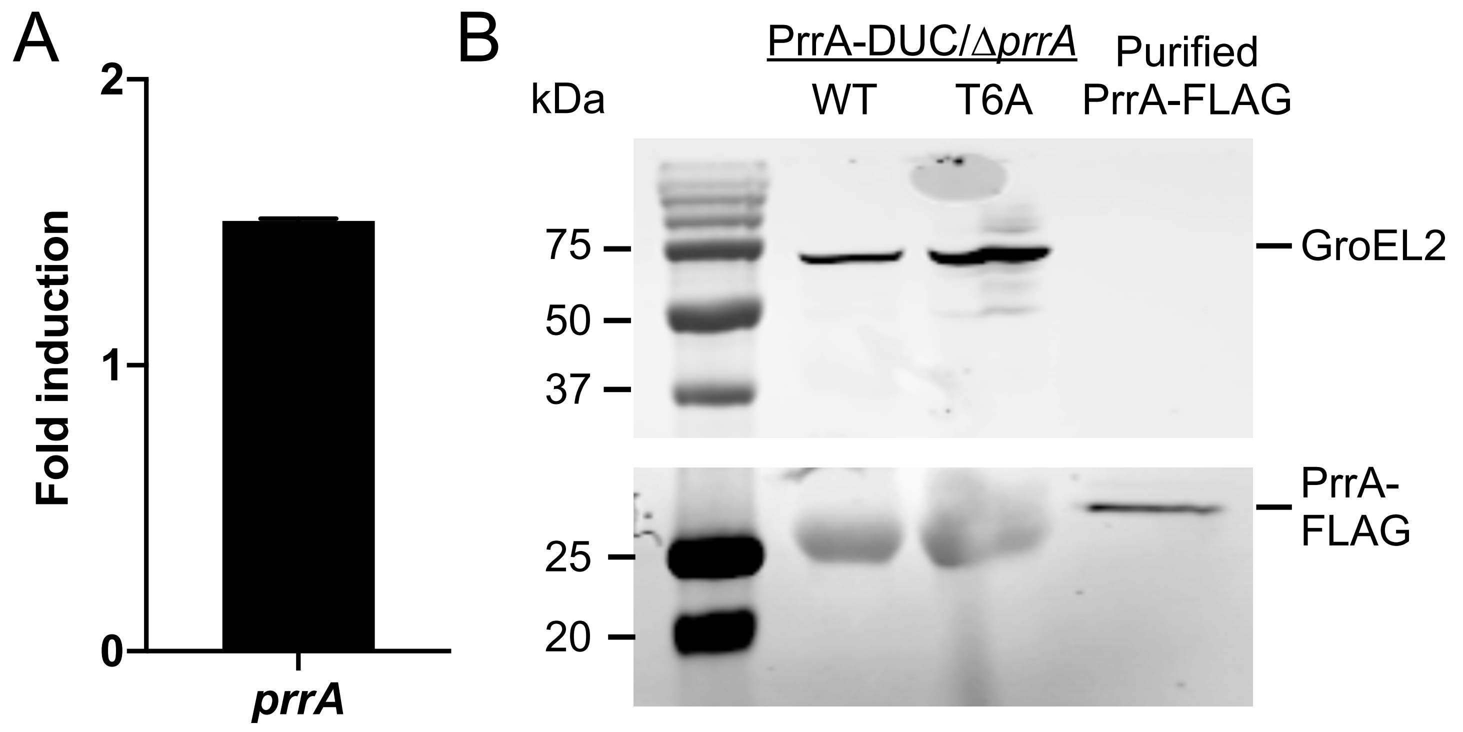


**Figure S6. Transcript and protein levels of PrrA are similar in the PrrA-DUC/Δ*prrA* and PrrA-T6A-DUC/Δ*prrA* strains.** (A) shows qRT-PCR of PrrA-DUC/Δ*prrA* and PrrA-T6A-DUC/Δ*prrA* Mtb grown in 7H9, pH 7.0 media for 4 hours. Fold induction compares *prrA* transcript levels in PrrA-T6A-DUC/Δ*prrA* to the PrrA-DUC/Δ*prrA* strain. *sigA* was used as the control gene, and data are shown as means ± SD from 3 technical replicates. (B) shows western blot analysis of PrrA-DUC/Δ*prrA* ("WT") and PrrA-T6A-DUC/Δ*prrA* ("T6A") grown in 7H9 pH, 7.0 media for 9 days, before culture was normalized to the lowest $OD_{600}$ and lysates prepared. Membranes were blotted with either an anti-GroEL2 antibody as a loading control (top panel) or an anti-FLAG antibody (bottom panel).

### Figure S7

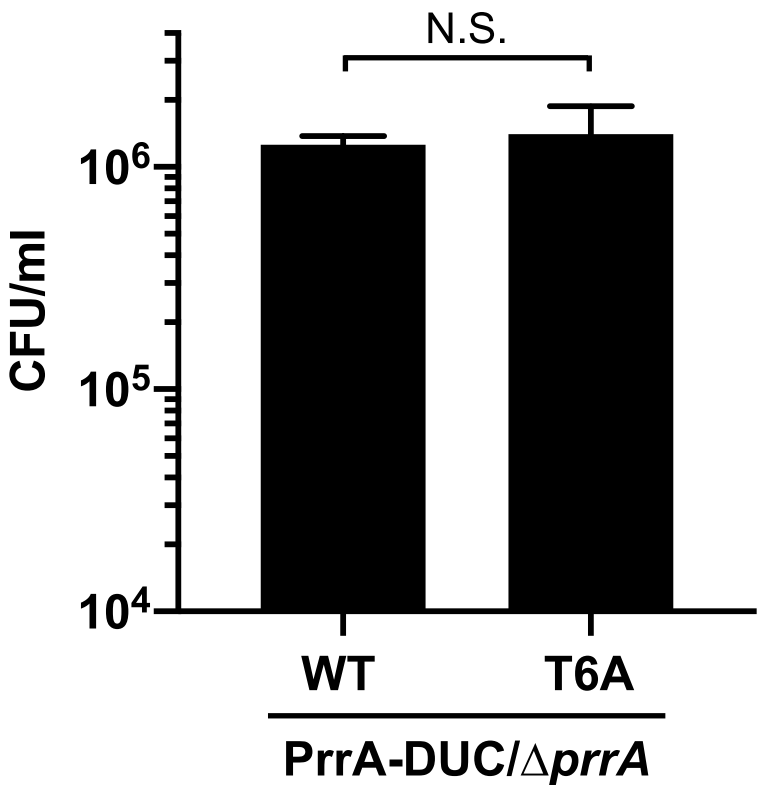


**Figure S7. Similar inoculum of Mtb strains used in the macrophage infection.** PrrA-DUC/Δ*prrA* ("WT") and PrrA-T6A-DUC/Δ*prrA* ("T6A") Mtb input preparations for macrophage infection were plated for colony forming units (CFUs) determination. Data are shown as means ± SD from three experiments. p-value was obtained with an unpaired t-test. N.S. not significant.
